# Learning Your Heart Actions From Pulse: ECG Waveform Reconstruction From PPG

**DOI:** 10.1101/815258

**Authors:** Qiang Zhu, Xin Tian, Chau-Wai Wong, Min Wu

**Affiliations:** Department of Electrical and Computer Engineering, University of Maryland, College Park, MD, 20742 USA, where the work was carried out, and is now with Facebook Inc.; Department of Electrical and Computer Engineering, University of Maryland, College Park, MD, 20742 USA; Department of Electrical and Computer Engineering, North Carolina State University, Raleigh, NC, 27695 USA

**Keywords:** ECG, PPG, inverse problem, DCT

## Abstract

This paper studies the relation between electrocardiogram (ECG) and photoplethysmogram (PPG) and infers the waveform of ECG via the PPG signals that can be obtained from affordable wearable Internet-of-Things (IoT) devices for mobile health. In order to address this inverse problem, a transform is proposed to map the discrete cosine transform (DCT) coefficients of each PPG cycle to those of the corresponding ECG cycle based on our proposed cardiovascular signal model. The proposed method is evaluated with different morphologies of the PPG and ECG signals on three benchmark datasets with a variety of combinations of age, weight, and health conditions using different training setups. Experimental results show that the proposed method can achieve a high prediction accuracy greater than 0.92 in averaged correlation for each dataset when the model is trained subject-wise. With a signal processing and learning system that is designed synergistically, we are able to reconstruct ECG signals by exploiting the relation of these two types of cardiovascular measurement. The reconstruction capability of the proposed method can enable low-cost ECG screening from affordable wearable IoT devices for continuous and long-term monitoring. This work may open up a new research direction to transfer the understanding of the clinical ECG knowledge base to build a knowledge base for PPG and data from wearable devices.

## I. Introduction

**C**ARDIOVASCULAR disease (CVD) has become the leading cause of human death – about 32% of all deaths worldwide in 2019 according to the Global Burden of Disease results [2]. Statistics also reveal that young people, especially athletes, are more prone to sudden cardiac arrests than previous generations [3]. Those life-threatening CVDs often happen outside clinics and hospitals, and the patients are recommended by cardiologists to attend a long-term continuous monitoring program [4].

The electrocardiogram (ECG) is a fundamental tool of clinical practice and today’s most commonly used cardiovascular diagnostic procedure [5]. Many modern wearable ECG systems have become simpler in physical configuration, more reliable than before, and may weigh only a fraction of a pound. However, despite the form-factor improvement in newer devices such as the Zio patch, the prolonged use of self-adhesive sensors, say, in a multi-day monitoring session, may increase the risk for skin irritations especially for persons with sensitive skins [6], [7]. Apple Watch can measure ECG signal by asking the user to tap its crown using a fingertip, but the requirement of active user participation precludes it from being used for long-term monitoring. To facilitate longterm ECG monitoring for a much broader range of users, it is imperative that the ECG edge sensor can operate without the need for user’s active participation.

The photoplethysmogram (PPG) is a noninvasive circulatory signal related to the pulsatile volume of blood in the tissue [8]. In common PPG modalities, the tissue is irradiated by a light-emitting diode, and the reflected or transmitted light intensity is measured by a photodetector on the same or the other side of the tissue. A pulse of blood modulates the light intensity at the photodetector, and the PPG is varying in an opposite direction with the volume of blood [8]. Compared with ECG, PPG is easier to set up and more economical, making it nearly ubiquitous in clinics and hospitals in the form of finger/toe clips and oximeters. In consumer electronics, PPG has increasing popularity in the form of wearable Internet-of-Things (IoT) devices that offer continuous and long-term monitoring capability with little to no skin irritations.

The PPG and ECG signals are intrinsically related, considering that the variation of the peripheral blood volume is directly influenced by the myocardial activities, and these activities are controlled by the electrical signals originating from the sinoatrial (SA) node. The timing, amplitude, and shape characteristics of the PPG waveform contain not only information about the connective vasculature but also rich information about heart activities. These features have been translated to measure heart rate, heart rate variability, respiration rate [9], blood oxygen saturation [10], blood pressure [11], and to assess vascular function [12]. As the prevailing use of wearable devices capturing users’ daily PPG signals, we are inspired to utilize this cardiovascular relation to reconstruct the ECG waveform from the PPG measurement. This exploration, if successful, can provide low-cost ECG screening for continuous and long-term monitoring and take advantage of both the rich clinical knowledge base of ECG signal and the easy accessibility of the PPG signal.

Regarding related art, there are papers on physiological parameter estimation for ECG or PPG, e.g., [13]–[15]. These works take as input time-series signals and output a few numbers/parameters that characterize the physiological status. There are also papers formulating binary decision problems for input ECG or PPG signals, e.g., [16]–[19]. Most of these works focused on the detection of rhythm-based disorder from the PPG signal rather than morphological disorder, and none of them directly estimates the ECG time-series signal from the PPG signal. A machine learning approach is introduced in [20] to train several classifiers to estimate ECG interval parameters from selected features of PPG. Even though the system achieved 90% accuracy on a benchmark dataset, only being able to estimate some parameters of ECG may limit the direct deployment of the method for ECG-based screening and monitoring. The ECG signal, rather than the derived parameters from a white-box or black-box system, is what intrigues some cardiologists since it allows them to leverage their medical expertise and clinical experiences for diagnosis.

In this paper, we propose to estimate the waveform of the ECG signal using PPG measurement by learning a signal model that relates the two time series. As a proof-of-concept for this line of research, we develop a principled learning framework and establish the baseline performance. More specifically, we introduce an interpretable inference framework motivated by the biophysical process of heartbeat cycle and signal-and-system modeling. Our initial investigation focuses on the following research questions and data science scenarios of PPG sensing data in near-ideal experimental conditions:

- Scenario 1 (*Subject-Specific Model*): Can the proposed learning system, trained by an individual’s signal pairs of ECG and PPG, predict his/her ECG waveforms from unseen PPG measurements?
- Scenario 2 (*Group Model*): Can a single model, trained by a group of subjects with certain trait combinations (e.g., age, weight), predict ECG waveforms from unseen PPG measurements for an individual in the training group?

Specifically, we first preprocess the ECG and PPG signal pairs to obtain temporally aligned and normalized sets of signals. We then segment the signals into pairs of cycles and train a linear transform that maps the discrete cosine transform (DCT) coefficients of the PPG cycle to those of the corresponding ECG cycle. The ECG waveform is then obtained via the inverse DCT. We evaluate our methodology on two publicly available datasets as well as a self-collected dataset, which in total contains 147 subjects with a wide variety of age, weight, and health conditions. As an example, Fig. 1 shows a five-second reconstructed ECG signal in the test set using the proposed method. Note that the reconstructed ECG signal is almost identical to the reference one.

**Fig. 1.**
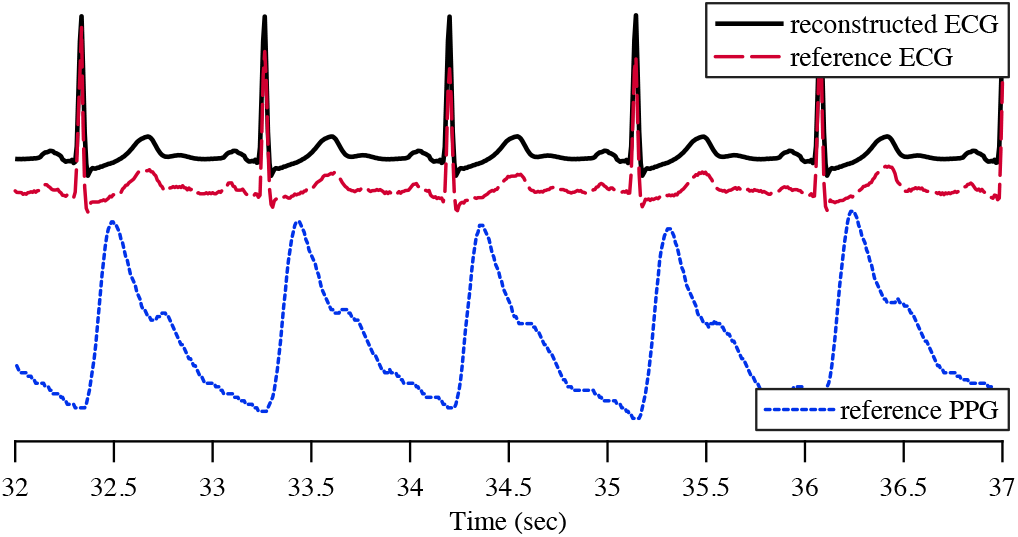
Upper: a five-second reconstructed ECG signal in the test set (black line) vs. the reference ECG signal (red line) using the data from the MIMICIII database [21]. The two signals were intentionally drawn with an offset in the vertical direction to better reveal the details. Lower: the corresponding PPG signal used to reconstruct the ECG signal.

Building upon and extending from the preliminary results reported in [1], the contribution of this work is fourfold. First, a physiological model for mathematically characterizing the relationship between the ECG and PPG time-series signals has been proposed and justified using electrical, biomechanical, and optophysiological principles. The proposed model provides an in-depth understanding of the two cardiovascular measurements and provides a principle for algorithm development along this line. Second, a principled learning framework based on the proposed physiological model is developed. The experiments using various datasets with a total 149 subjects suggest that the developed learning system can accurately reconstruct ECG signals from PPG signals when the system is trained in both subject-dependent and subject-independent manner. Third, to the best of our knowledge, together with our short conference paper reporting the promising preliminary results, this paper presents the first systematic framework addressing the problem of reconstructing ECG signals from the PPG signals. This research suggests an encouraging potential for a more user-friendly, low-cost, continuous, and long-term cardiac monitoring that supports and promotes public health, especially for people with special needs. This work may open up a new direction for cardiac medical practitioners, wearable technologists, and data scientists to leverage a rich body of clinical ECG knowledge and transfer the understanding to build a knowledge base for PPG and other data from wearable devices. Lastly, we extend the discussion of the system performance when the ground truth cardiac cycle information in the test set is unavailable and is estimated from the PPG signals. We present possible variations and extensions of our proposed system to facilitate the subsequent development of related art. We also discuss the possible scenarios when a subject is with certain cardiac implications or the PPG signal is significantly corrupted by the motion artifacts and requires proper denoising.

The rest of the paper is organized as follows. In Section II, we mathematically model the relationship between the ECG and the PPG signals. Based on the signal model, we introduce the proposed learning system in Section III. We evaluate it and report the experimental results in Section IV, and discuss possible extensions and limitations of the proposed system in Section V. The conclusion is drawn in Section VI.

## II. A Cycle-wise Signal Model for PPG and ECG

In this section, we discuss a physiological model adopted in this paper to develop the proposed algorithm. As shown in Fig. 2, during each cardiac cycle, the atrioventricular (AV) node receives the electrical signals originating from the SA node. The AV node then transmits these bioelectric signals sequentially through His bundle, the left bundle branch, and Purkinje fibers to the left ventricular myocardium, causing the depolarization and contraction of the left ventricle. As a result of this process, the pressure of the left ventricle rises and exceeds the aortic pressure, causing the opening of the aortic valves, blood flow from the left ventricle into the aorta, and the corresponding rise of the aortic pressure. Upon closure of the aortic valves, the generated pulse wave transmits the blood to the peripheral parts of our body, such as fingertips or toes, through a network of blood vessels. During the modeling process to be explained below, we have taken into account the following four main physiological factors: 1) human body resistance of the electrical path from the heart to the skin surface, 2) the myocardial activities that originate the aortic pressure wave, 3) the blood vessel as a channel from aortic pressure to the peripheral pulse wave, and 4) the transmissive/reflective strength of the skin tissue.

**Fig. 2.**
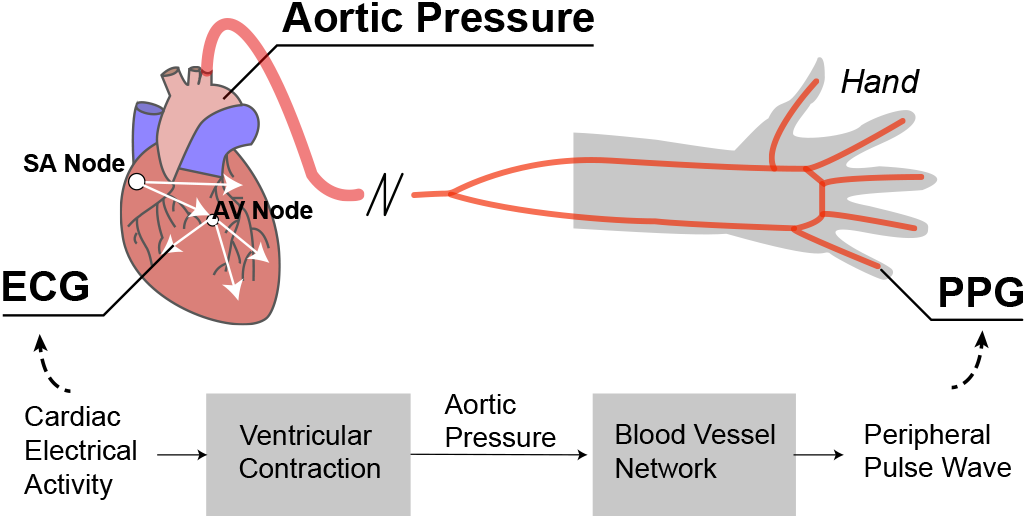
A visualization of the relationship between the ECG, the aortic pressure, and the PPG.

### A. The ECG Signal and the Aortic Pressure

Consider one specific cardiac cycle. We denote the uniformly sampled cardiac electrical activity as *e*(*n*), *n* ∈ [1, *L*], where *L* is the total number of samples within the cycle. We denote the electrocardiogram measurement recording the potential difference between two electrodes placed on the surface of the skin as *c_y_*(*n*). Taking into account the human body electrical resistance and the sensor noise, we model the ECG signal *c_y_*(*n*) as:

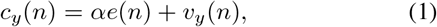

where *α* denotes a subject-specific parameter accounting for the resistance of the electrical path between the heart and the skin surface; *v_y_*(*n*) denotes the ECG sensor noise, which is modeled as a zero-mean white Gaussian process.

The contraction and relaxation of the heart muscles follow the bioelectric activities of the heart. These biomechanical activities further modulate the aortic pressure via the opening and closing of the aortic valves. The aortic pressure, denoted as *p_a_*(*n*), is therefore caused by the cardiac electrical activities *e*(*n*). In our proof-of-concept work, we propose to capture the dominant relationship between *p_a_*(*n*) and *e*(*n*) using linear mappings between their corresponding coefficients of the DCT representation [22] shown as follows:

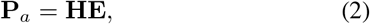

where **E**, 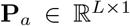 are the vectors of the DCT-II coefficients of *e*(*n*) and the aortic pressure *p_a_*(*n*), respectively. Transition matrix 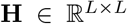 accounts for the myocardial activities. It has been shown in the literature that DCT can provide a representation that is compact [23], which allows us to build models that can capture relationships more efficiently. Note that the relation between the two signals **E** and **P**_*a*_ is expected to be not as simple. The linear model is introduced to approximate and reveal insights of this complex physiological system up to the first order.

### B. The Pulse Wave and the PPG Signal

When the pulse wave and blood flow travel through our body from the aorta to a peripheral site, it might experience different interactions with the blood vessels, for instance, splitting and pushing. We assume the structure of the blood vessel path of a specific person is time-invariant. Inspired by models for the vocal tract in speech production ( [24], Chapter 3), we propose to model this blood vessel channel from the aorta to the peripheral site as a linear time-invariant system. We denote the peripheral pulse signal at a specific body site as *p_p_*(*n*). We write *p_p_*(*n*) according to the prior channel assumption as:

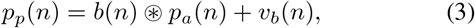

where *b*(*n*) denotes the impulse response of the channel of blood vessels, and ⊛ denotes a symmetric convolution operation [22]. *v_b_*(*n*) is the zero-mean white Gaussian noise, capturing the variance of this model. The symmetric convolution of *b*(*n*) and *p_a_*(*n*) gives a result that is the same as a linear convolution of the symmetrically left-sided extended version of *b*(*n*) and two-sided extended version of *p_a_*(*n*). The extension of *p_a_*(*n*) provides smooth boundary values for filtering near its original endpoints. This “folded aliasing” may be preferable in modeling this blood vessel channel effect to the wrap-around aliasing of a circular convolution [22].

We assume the PPG sensor attached to the same peripheral site works in the transmissive setup. It means that the photodetector of the PPG sensor is on the other side of the tissue with the light-emitting diode. We assume the light source has a constant intensity of *I* on the spectral range of the receiver side. We further assume no relative motion between the attached skin and the photodetector, and the contact is tight enough so that the signal is not influenced by the possible environmental illuminations. We write the PPG measurement, denoted as *c_x_*(*n*), as:

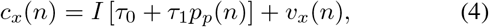

where *τ*_0_ and *τ*_1_ denote the relative transmissive strength of the non-pulsatile components and pulsatile components of tissue, respectively [25].^1^ *v_x_*(*n*) denotes the PPG sensor noise, which is modeled as a zero-mean white Gaussian process. We can rewrite (4) as:

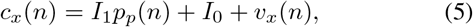

where *I*_1_ = *Iτ*_1_ and *I*_0_ = *Iτ*_0_.

### C. The Inverse Model from PPG to ECG

According to the property of the symmetric convolution, the type-II DCT of a symmetric convolution of two signals is the element-wise product of type-II DCT of one signal and type-I DCT of the other signal [22]. Combined with the linearity property of the DCT, we can rewrite (3) in the frequency domain as:

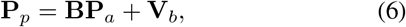

where **P**_*p*_, **P**_*a*_, and 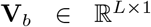 are the DCT-II coefficients of *p_p_*(*n*), *p_a_*(*n*), and *v_b_*(*n*) respectively. 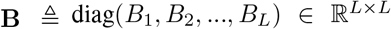, where *B_k_* denotes the *k*th DCT-I coefficient of *b*(*n*). We next apply a type II DCT on both sides of (1) and (5) and we arrive at:

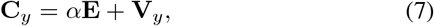

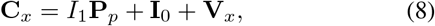

where **C**_*y*_, **V**_*y*_, **C**_*x*_, **I**_0_ and 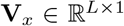 denotes the DCT-II coefficients of *c_y_*(*n*), *v_y_*(*n*), *c_x_*(*n*), constant function *I*_0_ and *v_x_*(*n*) respectively. Assuming the nonsingularity of the matrix **B** and **H** and according to (2), (6), (7), and (8), we have:

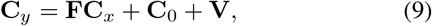

where 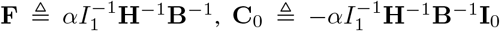, and 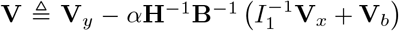. When we look individually at each element of **C**_*y*_, we have:

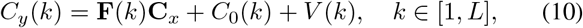

where **F**(*k*) is the *k*th row of matrix **F**; *C*_0_(*k*) and *V*(*k*) denote the *k*th element of **C**_0_ and **V**, respectively. We know *V*(*k*) is a zero-mean Gaussian random variable, as it is a linear combination of zero-mean Gaussian random variables from *v_y_, v_b_*, and *v_x_*. According to (10), the relation between the PPG and the ECG signal is captured by a linear model in their frequency domain. We are thus motivated to explore the linear relationships between the DCT coefficients of PPG signal and those of the ECG signals.

## III. Proposed Method for PPG to ECG Conversion

Due to the influence of various physiological factors discussed in Section II, the generation of the PPG signal from the ECG signal via successive steps can be regarded as a lossy process from the signal processing viewpoint. For example, if we regard the blood vessels as a lowpass system, the high-frequency information of the aortic pressure wave is lost when the blood reaches the peripheral sites. We are motivated by such established inverse problems as image deblurring [27] and audio bandwidth extension [28] to exploit the signals’ inherent learnable correlation among components of different frequencies. Although the high-frequency portion of the spectrum from the upstream ECG signal suppressed by the equivalent lowpass filter and would not have been recoverable by a linear, time-invariant filtering process, the high-frequency spectrum can be inferred from other spectral components of the PPG signal through a well-designed model and learning methodology. According to the signal model we discussed in the previous section, we propose a system that learns the linear transform **F** from pairs of PPG and ECG data. The pipeline of the system is shown in Fig. 3. The pair of PPG and ECG signals are first preprocessed into synchronized cycles. The cycle pairs are then fed into the training system to learn the transform matrix. We discuss further the details of the system as follows.

**Fig. 3.**
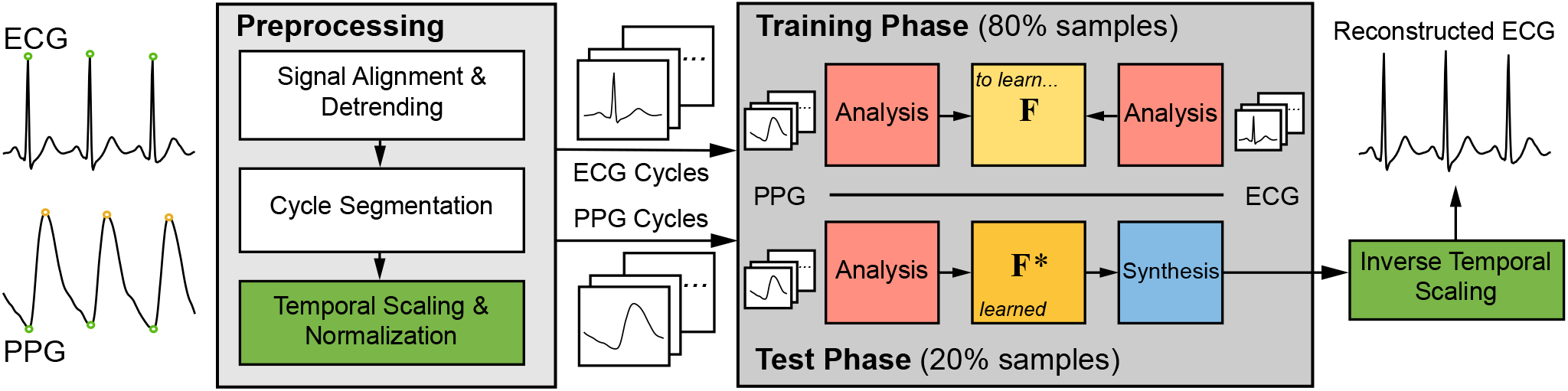
Flowchart of the proposed system. The ECG and PPG signals are first preprocessed to obtain physically aligned and normalized pairs of cycles. The selected DCT coefficients of 80% pairs of cycles are used for training a linear transform **F** which is used in the test phase to reconstruct the ECG signals.

### A. Preprocessing: Cycle-wise Segmentation

We first preprocessed the ECG and PPG signals to obtain temporally aligned and normalized pairs of the signal cycles to facilitate the investigation in the subsequent training stage. The left part of Fig. 3 shows the preprocessing phase, which includes data alignment, signal detrending, cycle-wise segmentation, temporal scaling, and normalization stages that be explained as follows.

#### a) Data Alignment and Detrending

As the relation between ECG and PPG is modeled at the cycle level, it is crucial first to estimate the signal delay in each trail and temporally align the signals. To achieve this goal, we propose a two-level signal alignment scheme. We first use the peak features of the signal pair to estimate the cycle-wise delay. We then align the signal to the sample level based on the physical meaning and correspondence of the two signals.

Suppose we have a pair of almost simultaneously recorded PPG and ECG signals, denoted as 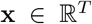 and 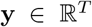 respectively. We name the coordinate of the systolic peak in the *i*th cycle of PPG as *n*_sp_(*i*) and the R peak of ECG as *n*_rp_(*i*). The cycle delay *m*_delay_ is searched for in a discrete interval 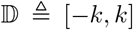. For each evaluated 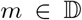, we first preliminarily align the signal with respect to 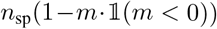, and 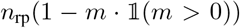. The aligned coordinates of PPG and ECG peaks are 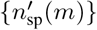 and 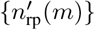. The cycle delay 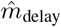 is estimated by solving the following problem:

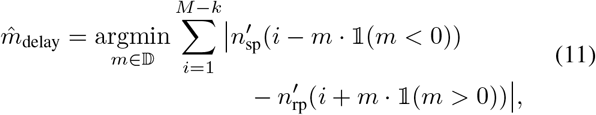

where *M* denotes the total number of cycles; 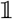 denotes the indicator function. After we estimate the cycle delay 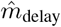, the PPG signals are shifted so that the systolic peaks of PPG and the R peaks of ECG are temporally matched.

We then align the R peak of the ECG and the onset point of PPG in the same cycle, considering that the R peak corresponds approximately to the opening of the aortic valve, and the onset point of PPG indicates the arrival of the pulse wave [8]. In this way, the PPG and ECG signals are aligned within the cycle according to their physiological correspondence.

The quasi-DC components in both signals caused by respiration, vasomotor activity, and thermoregulation [29] require additional attention to temporal pattern analysis. With the prior information that such nonstationarities represent slowly-varying trends in the signal, we estimate the trends from the s using the *smooth prior method* introduced in [30] and subtract the trends from the original signals.

#### b) Segmentation & Normalization

We next segment each cycle of the signal 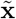 and 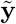 to prepare for the learning phase. In this paper, two cycle-segmentation schemes called *SR* and *R*2*R* are compared. They are specified as follows.

- *SR*: we segment the signal according to the points which are 1/3 of the cycle length to the left of the R peaks of the ECG signal. We call this scheme SR as it approximately captures the standard shape of sinus rhythm.
- *R2R*: we segment the signal according to the location of the R peak of the ECG signal to mitigate the reconstruction error in the QRS complex.

After the segmentation, each cycle sample is scaled in time and amplitude to make it of equal length *L*, zero mean, and unit sample standard deviation. We denote the normalized PPG and ECG cycle samples as **c**_*x*_, 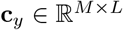.

### B. Learning a DCT-domain Linear Transform

The right part of Fig. 3 shows our proposed learning framework. The linear transform **F** in the signal model (10) is learned in the training phase and is used to reconstruct the ECG waveform in the test phase.

Specifically, we first perform cycle-wise DCT on **c**_*x*_ and **c**_*y*_, which yields **C**_*x*_, 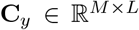. Then the first *L_x_, L_y_* DCT coefficients of **C**_*x*_, **C**_*y*_ are selected to represent the corresponding waveform as the signal energy is concentrated mostly on the lower frequency components per our observation. We denote them as 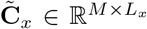 and 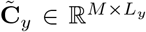. We next separate 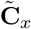 and 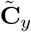 into training and test sets as 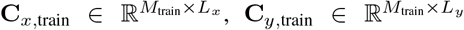 and 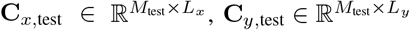, where *M*_train_ + *M*_test_ = *M*.

In the training process, we propose to learn the linear transform matrix 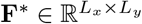 that maps from PPG to ECG DCT coefficients using three least square methods. They are ordinary Least Square (oLS), Ridge regression (Ridge), and Least Absolute Shrinkage and Selection Operator (Lasso).

The oLS solution of **F** is the minimizer of residue sum-of-squares of the ECG DCT coefficients:

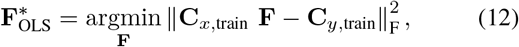

where ║·║_F_ denotes the Frobenius norm of a matrix. The oLS generates the most straightforward closed-form solution 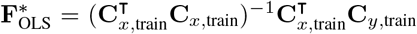 with low prediction bias, but its estimates often have large prediction variance. Prediction accuracy can sometimes be improved by regularized least square methods, such as the Ridge and Lasso.

The Ridge adds a regularization term after the OLS formulation to shrink the size of **F**. The Ridge estimate is defined by

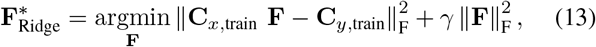

where *γ* > 0 is a complexity parameter that controls the shrinkage of **F** toward zero thereby reducing the variance of the predictions [31]. The analytic solution to (13) is 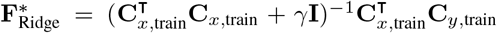, where **I** is the identity matrix.

The Lasso is another shrinkage method similar to Ridge, but replaces the penalty 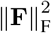 with ║**F**║_**1**_. Note that this subtle difference may eventually lead to a completely different solution with the “soft thresholding” of the entries in 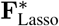, and thus give high interpretability of the model. We solve the Lasso with the *alternating direction method of multipliers*, which is introduced in [32].

In the test phase, we apply the optimal linear transform **F**\* learned in the training stage on **C**_x,test_ and estimate the corresponding DCT coefficients of ECG cycles. We denote the estimate as 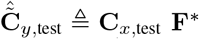. To reconstruct ECG, we first augment each row of 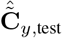 to be in the same dimension as L (by padding zeros). We denote the zero-padded matrix as 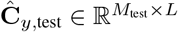. We then apply inverse DCT to each row of 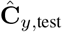, interpolate the resulted matrix row by row to its original temporal scale, and concatenate the inversely scaled pieces of cycles to obtain the reconstructed ECG signal **ŷ**_tes_t.

## IV. Experiments

### A. Experiment I: Capnobase TBME-RR Database

We first used the Capnobase TBME-RR [9] to evaluate the performance of the proposed system. The dataset contains 42 eight-min sessions of simultaneously recorded PPG and ECG measurements from 29 pediatric and 13 adults ^2^, sampled at 300 Hz. The 42 cases were randomly selected from a larger collection of physiological signals collected during elective surgery and routine anesthesia. Each recorded session corresponds to a unique subject. The PPG signal was acquired on the subjects’ fingertips via a pulse oximeter. The dataset has a wide variety of patient’s age (min: 1, max: 63, median: 14) and weight (min: 9 kg, max: 145 kg, median: 49 kg) and is thus a favorable dataset for testing the performance of our proposed system.

We first pruned the signals according to the human-labeled artifact segments and processed the pairs of ECG and PPG signals using the method introduced in Section III-A to obtain aligned and normalized pairs of the signal cycles. We set *L* = 300 and *L_y_* = 100, as most of the diagnostic information of ECG is contained below 100 Hz [5]. We set *λ* = 500 and *γ* =10 empirically as they offer the best regularization results in the tasks. In order to test the consistency of the system, we selected the first 80% of the data from each subject as the training set and the rest for testing. We used the following two metrics to evaluate the system performance in the test set:

- Relative root mean squared error:

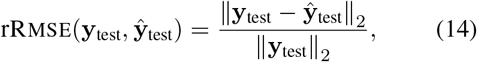
- Pearson’s correlation coefficient:

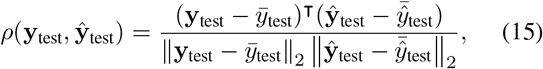

where **y**_test_, 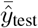, and 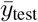 denote the ECG signal in test set, the average of all coordinates of the vectors **ŷ**_test_ and **y**_test_ respectively.

In this study, we evaluate the system in the following two training setups:

- *Group Model (GM)*: We trained a single linear transform matrix **F**^*^ using the training data from all subjects, i.e., we use a generic model to capture the signal relation for a group of subjects.
- *Subject-Specific Model (SM)*: A linear transform matrix **F**^*^ was trained and tested for each individual to obtain a subject-specific model.

We first cross-validated the number of DCT coefficients of the PPG signal, *L_x_*, used in the learning system. It is clear that the more variables as predictors, i.e., more PPG DCT coefficients are used in the linear system, the better the performance can be achieved in training. However, we can observe from Fig. 4 that the performance of our system in the test set using either SR and R2R becomes saturated as *L_x_* grows to approximately 18 and 12 in the group model and subject-specific model setup, respectively. The trends of convergence in both setups suggest potential model overfitting. Another observation is that the convergence rate is slower in the group model setup compared with the subject-specific model setup. Such observation is expected because the data diversity is much higher in the group model setup than that in the subject-specific model setup, and more variables are needed to capture the additional variance in the group model setup. *L_x_* = 18 in the group model setup and *L_x_* = 12 in the subject-specific model setup are thus favorable to us as the system has comparable performance and the model is parsimonious than those with larger *L_x_*.

**Fig. 4.**
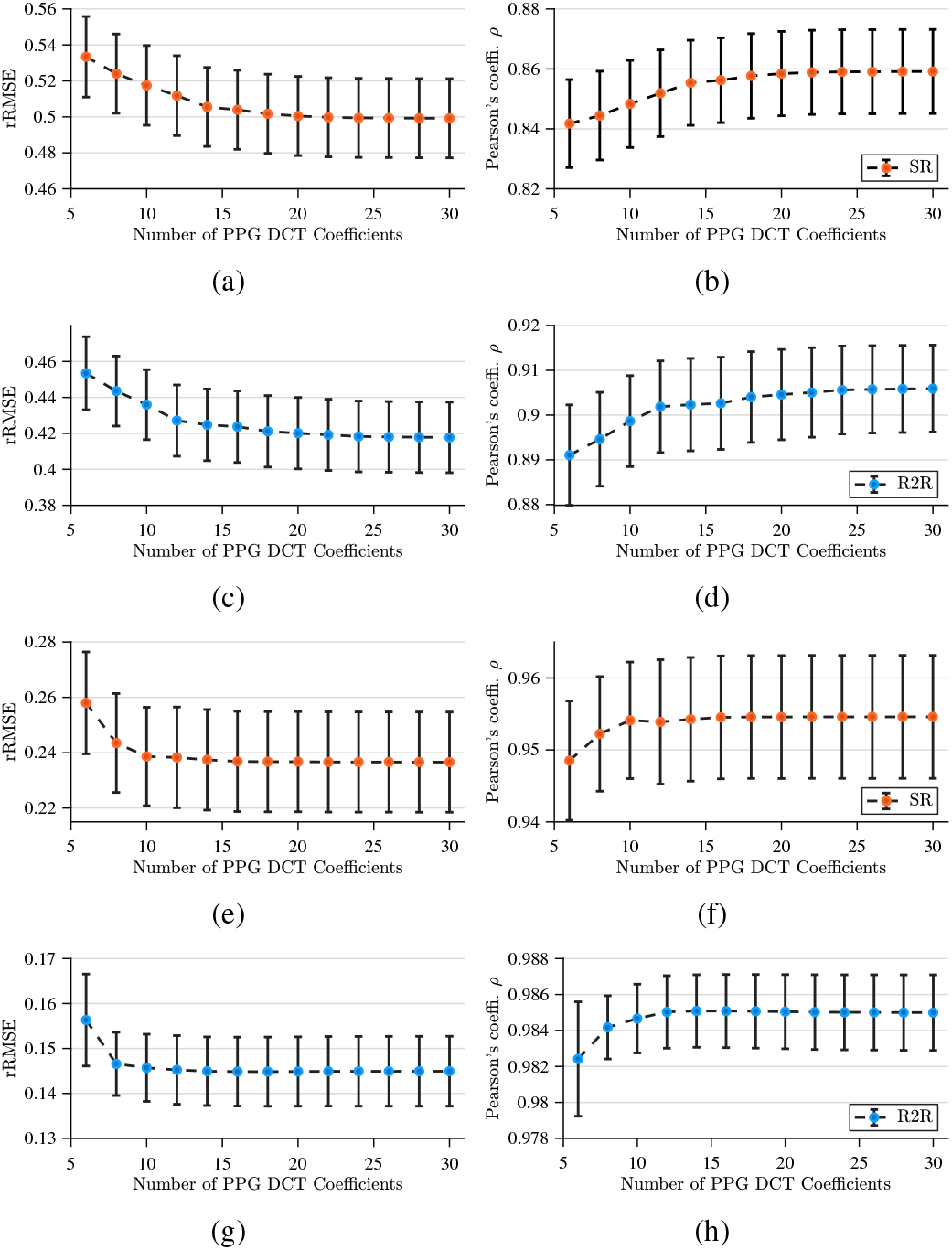
The line plots give the average of rRMSE in (a), (c), (e), and (g) and *ρ* in (b), (d), (f), and (h) of all sessions in the test set for different choices of the number of PPG DCT coefficient *m*_1_ using SR (a), (b), (e), and (h) and R2R (c), (d), (g), (h) segmentation scheme and group model (a)–(d) and subject-specific model (e)-(h) setup respectively. The vertical bar at each data point shows one standard error above and below the sample mean.

The norm of one cycle of ECG signal is usually dominated by that of the QRS complex. This fact of unbalanced signal energy distribution might lead to insufficient evaluation on the P wave and T wave of the ECG signal. To address this problem, we further separated the ECG cycle into and evaluated the system performance on segments of the P wave, QRS wave, and T wave. The evaluation was performed in terms of rRMSE and *ρ* on each segment as well as using the entire cycle of the signal. Specifically, we adopted the QRS detection algorithm introduced in [33] to locate the onset and endpoint of the QRS complex. We empirically selected the 60% point between the onset points of two adjacent QRS complexes as the separating point for the P and T waves. Fig. 5 shows one example of the ECG segmentation result sampled from the first subject in the database. Note that the onset and endpoint of all waves in each cycle are accurately estimated.

**Fig. 5.**
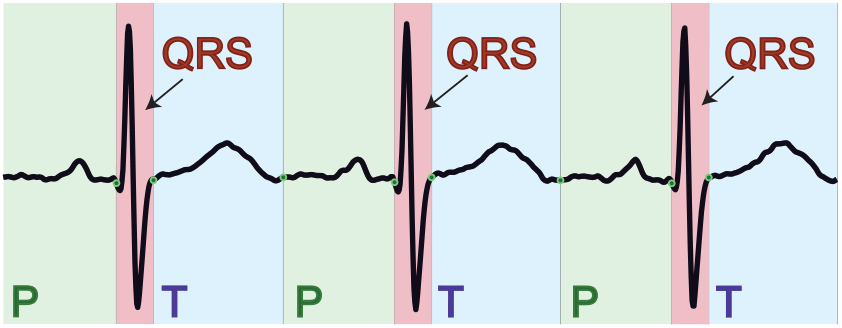
An example of the ECG segmentation result on three cycles of the signal in the 1st session of TBME-RR database. The green, red, and blue areas in the plot denote the estimated P waves, the QRS waves, and the T waves, respectively. For each cycle, the ratio between the duration of the QRS+T wave is 3/2 of the duration of the T wave.

We list the average performance using R2R and SR cycle segmentation schemes in different training setups in Table I and plot the results using the box plots in Fig. 6. In Table I, the performance is characterized by the sample mean and standard deviation of rRMSE and *ρ* on P, QRS, T, and all waves, where all wave denotes the whole length of the signal including every wave. From the statistics, we learn that overall R2R gives better performance than SR, and model trained in the subject-specific model setup gives better performance compared with that trained in the group model setup in this dataset as possible subject differences in terms of **H** in (2) and *b*(*t*) in (3) are expected. The three regression methods, OLS, Ridge, and Lasso give comparable performances. In general, R2R outputs comparable results on P and T waves compared with SR, whereas R2R outperforms SR on QRS and all waves. In the subject-specific model setup, the average performance in *ρ* on T wave is about 0.92 using R2R and 0.90 using SR, much higher values than those on the P wave. There are two possible reasons that explain this result. First, compared with the QRS and T waves, the amplitude of the P wave is much smaller. As a result, the P wave becomes more sensitive to the noise compared with the T wave. Second, the shape of the T wave signifies the repolarization of the ventricles, and the ventricular repolarization is correlated with the shape of the dicrotic notch in the PPG signal. This is because, during the ventricular repolarization process, the closure of the aortic valve is associated with a small backflow of blood into the ventricle and a characteristic notch in the aortic pressure tracings. This connection between the P wave of ECG and the dicrotic notch of PPG may facilitate the system performance on the P wave.

**Fig. 6.**
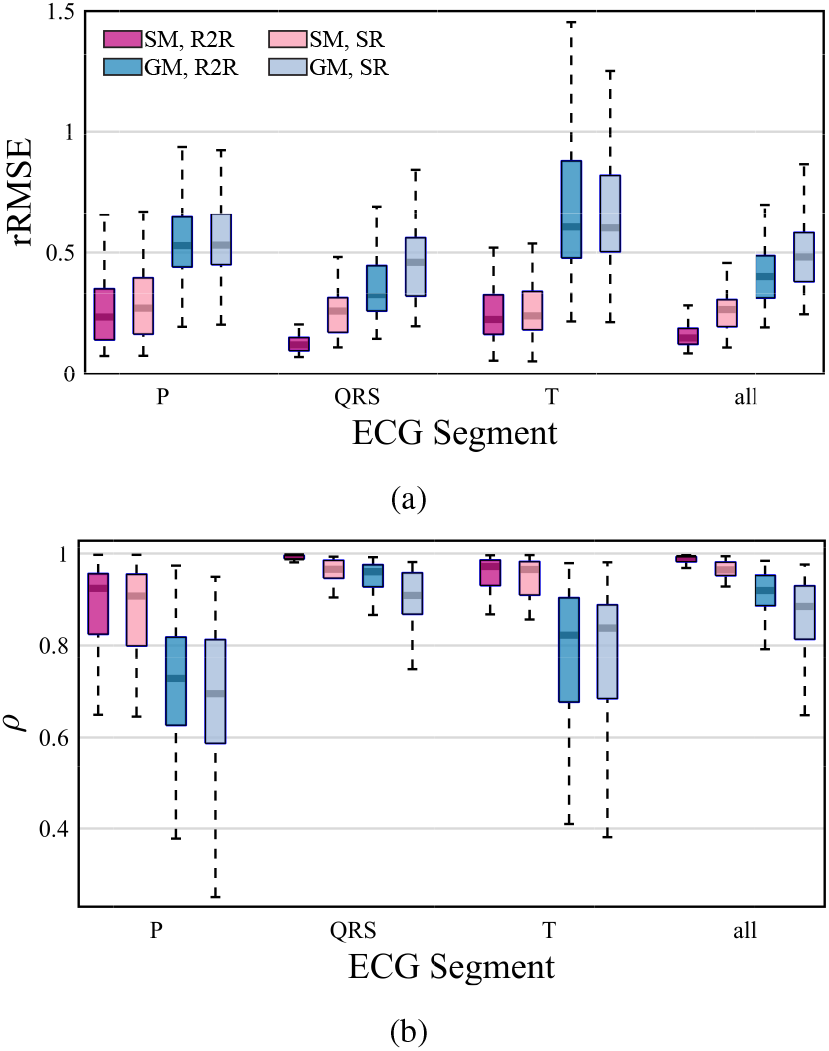
Comparison of the performance of the proposed method in test set of the TBME-RR database in different combinations of the SR or R2R segmentation schemes and the subject-specific model or group model test setups evaluated at P, QRS, T, and all waves. Statistics of the (a) rRMSE and (b) *ρ* are summarized using the box plots.

**TABLE I.**
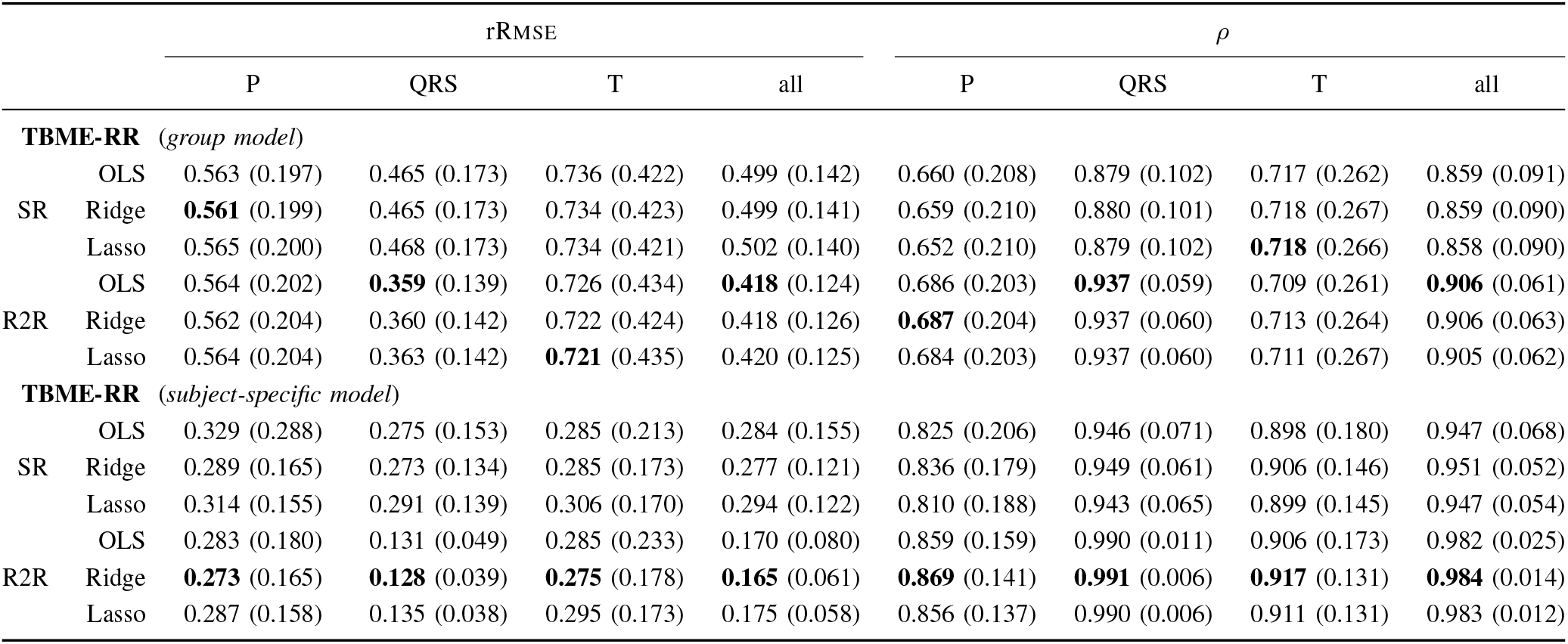
The system performance in test set of the TBME-RR database in terms of mean and standard deviation (in parenthesis) of RRMSE and *ρ*. R2R segmentation using different combinations of the training setup (GM/SM), the segmentation schemes (SR/R2R) and the linear regression methods (OLS/RIDGE/LASSO). The best performed entry in each column and training setup is bolded for better visualization. The entry with lowest standard deviation will be bolded if the means of multiple entries are identical.

As an example, we show a five-second segment of the reconstructed ECG waveform in the test set of the first subject in Fig. 7 using the R2R cycle segmentation scheme with *L_x_* = 18 in the group model setup and *L_x_* = 12 in the subject-specific model setup. We choose the first subject to be the example as the system performance evaluated on this subject approximates the average performance over the database. We see from the plot that the system retains most of the shape of the original ECG waveform except for the S peaks in the group model setup and almost perfectly reconstructs the ECG waveform and maintains the location of each PQRST peak in the subject-specific model setup.

**Fig. 7.**
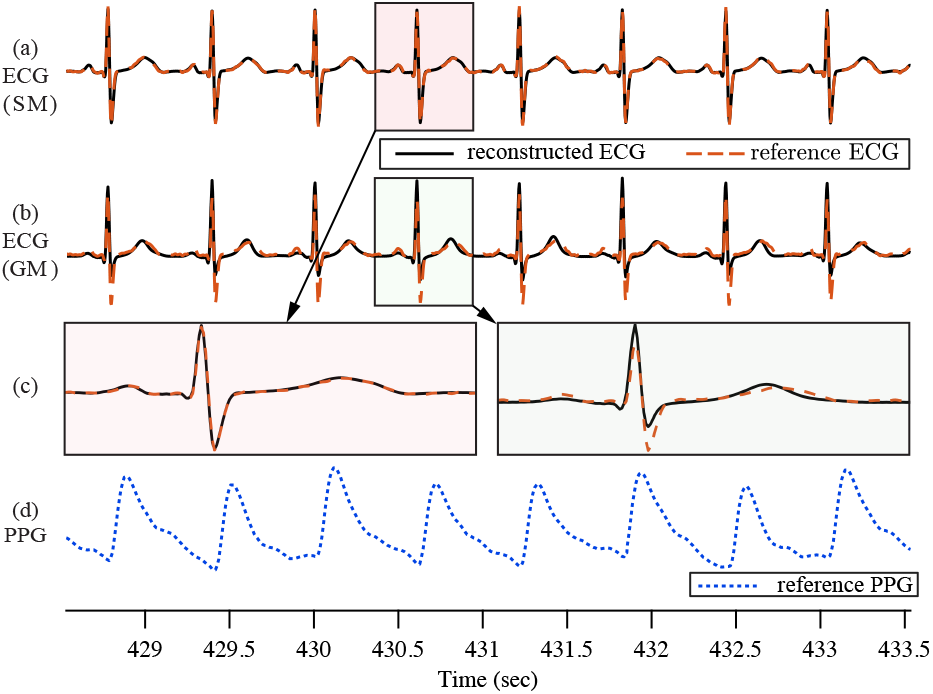
The reconstructed ECG (black solid line) in (a) the subject-specific model and (b) group model and the reference ECG (orange dashed line) waveform of the last 5 seconds of the first session (age: 4 years old, weight: 18 kg) in TBME-RR database. Zoomed-in version of the shaded cycle in each setup is shown in (c). The corresponding PPG waveform is shown in (d).

In Fig. 8, we plot the rRMSE and *ρ* of each session concerning subjects’ age and weight respectively in two 3-D plots in the group model and subject-specific model setup. We then fit a linear model with an interaction term for each combination of training setup and evaluation metric according to the least-squares criterion. An *F*-test is performed to test whether subjects’ profiles, i.e., age, weight, and the interaction between age and weight, can significantly affect the performance of the algorithm in each metric and training setup combination. *F*-test results of small *p*-values shown in Figs. 8(a) and 8(b) for the group model setup reveal that the performance of the algorithm is dependent on the combination of subject’s age and weight, whereas the large high *p*-values shown in Figs. 8(c) and 8(d) for the subject-specific model setup do not show strong evidence to reject the hypothesis that the performance of the algorithm is independent of age and weight. Moreover, we notice that the performance tends to be lower as the subject’s weight gets larger. This trend of performance degradation might be due to the bias of the training sample that the number of new-borns is much larger than the number of other groups of subjects in the database.

**Fig. 8.**
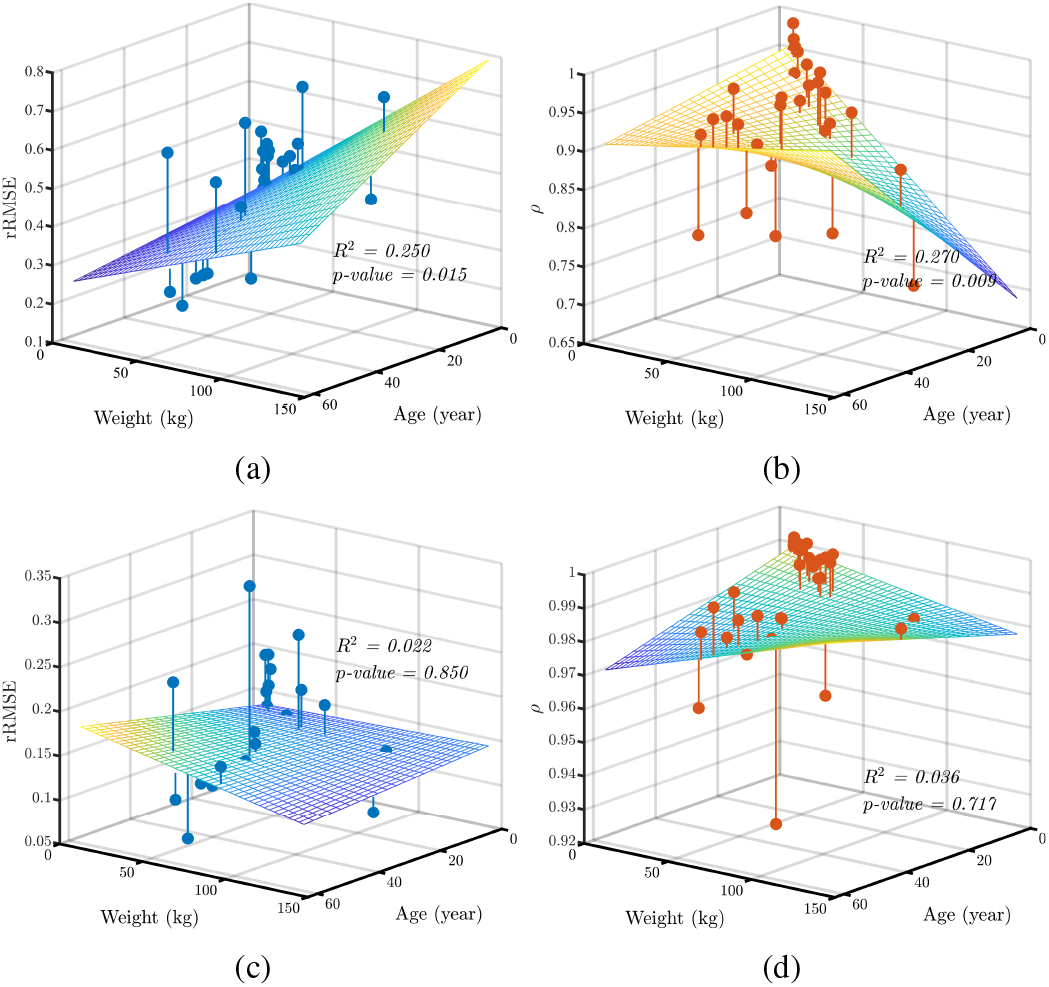
Scatter plots of (a) rRMSE and (b) *ρ* vs. subjects’ weight and age using R2R scheme. Each sample corresponds to one of 42 sessions. The surface mesh on each plot shows the regressed linear model: rRMSE or *ρ* ~ intercept + age + weight + age × weight. The *R*^2^ and the *p*-value of *F*-test is shown on each plot.

### B. Experiment 2: MIMIC-III Database

Medical Information Mart for Intensive Care III (MIMICIII) [21] is an extensive database comprising vital sign measurements at the bedside documented in the MIMIC-III waveform database and part of the patients’ profile in the MIMIC-III clinical database. The database is publicly available and encompasses a large population of ICU patients. In this experiment, a subset of the MIMIC-III database was used to evaluate the system’s performance when the subjects were with various cardiac or non-cardiac malfunctions.

Specifically, we selected waveforms that contain both lead II ECG and PPG signals from folder 35 in the MIMIC-III waveform database. Then we linked the selected waveforms with the MIMIC-III clinical database by subject ID to match with the corresponding patient profile. Among the patients, we selected those with specific cardiac/non-cardiac diseases and removed low signal quality PPG/ECG pairs. The resulting collected database consists of 53 patients with six common cardiac diseases and 50 patients with five types of noncardiac diseases. The distribution of the collected patients is visualized in stacked bar plots based on each one’s age group and disease type in Fig. 9. Each patient has three sessions of 5-min ECG and PPG recordings collected within several hours. Cardiac diseases in the resulting database include atrial fibrillation, myocardial infarction, cardiac arrest, congestive heart failure, hypotension, hypertension, and coronary artery disease, while non-cardiac diseases are composed of sepsis, pneumonia, gastrointestinal bleed, diabetic ketoacidosis, and altered mental status.^3^

**Fig. 9.**
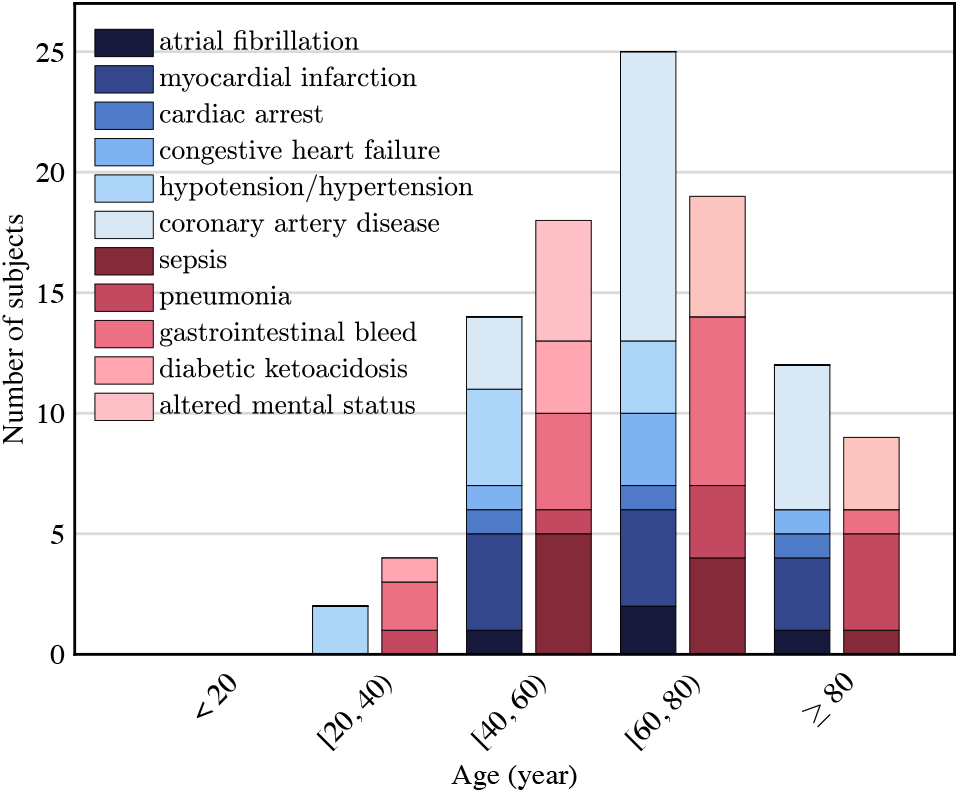
Distribution of subjects collected from the MIMIC-III database in five age groups and eleven disease types. Within each age group, the cardiac-related diseases are colored as different shades of blue on the left, and the noncardiac-related diseases are colored as different shades of red on the right.

In this part of the experiment, we evaluate our proposed system in the group model and subject-specific model setups (both under the R2R segmentation scheme). In group model setup, we trained one linear transform **F**\* using training data from patients with cardiac diseases and another linear transform **F**\* from non-cardiac disease patients, i.e., the trained model is independent with each subject in terms of disease type. In subject-specific model setup, we use the first two sessions for training and the third session for testing, i.e., to report the results for each individual.

We summarized the average performance in Table II using R2R cycle segmentation scheme in the subject-specific model and group model training setups. Results in Table II are characterized by the sample mean and the standard deviation of rRMSE and *ρ* values on P, QRS, T, and all waves as in the first experiment. The rRMSE and *ρ* values are also plotted using the box plots in Fig. 10. The statistics reveal that overall non-cardiac cases give better performance than cardiac cases as less variation exists in the morphology of non-cardiac ECG signals. The model trained in the subject-specific model setup gives better performance compared with that trained in the group model setup in this dataset, which suggests that **H** in (2) and *b*(*t*) in (3) may be subject-dependent. In general, for the subject-specific model setup, the average performance in *ρ* on T wave is about 0.90 and on QRS wave is about 0.94 using R2R, much higher than those on the P wave, which is in accordance with the first experiment.

**Fig. 10.**
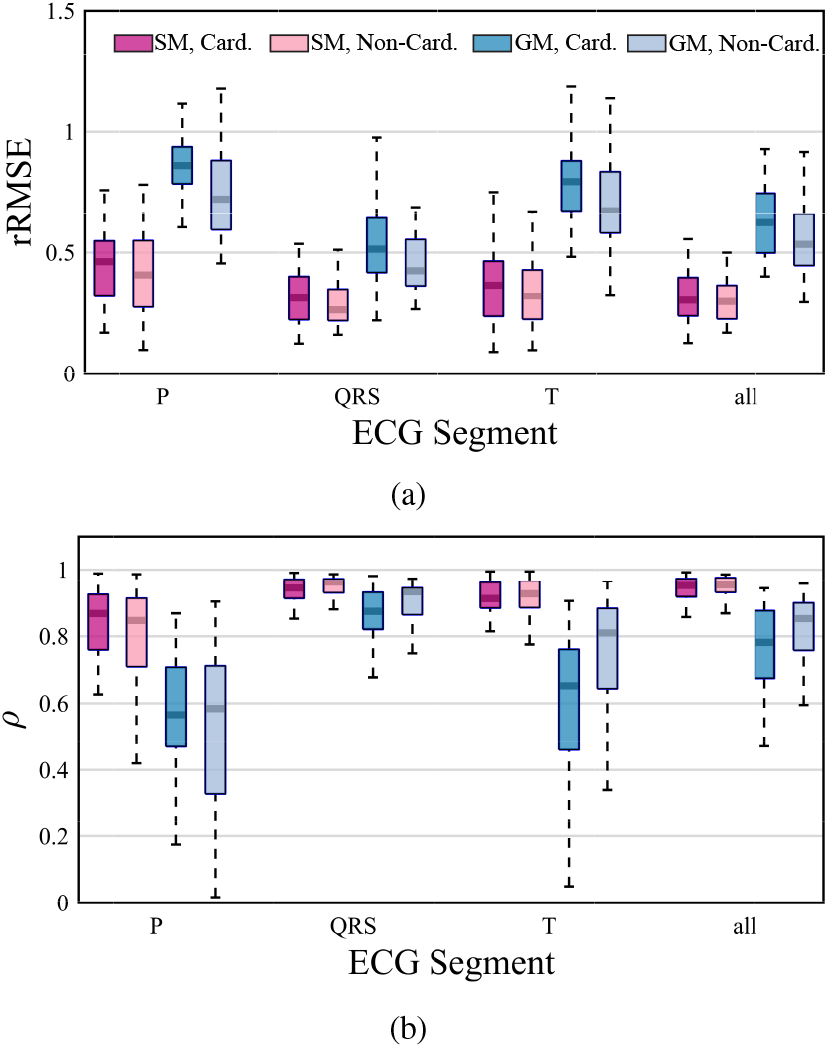
Comparison of the performance of the proposed method in test set of the MIMIC-III database in different combinations of the disease types and test setups. Statistics of the (a) rRMSE and (b) *ρ* are summarized using the box plots.

**Fig. 11.**
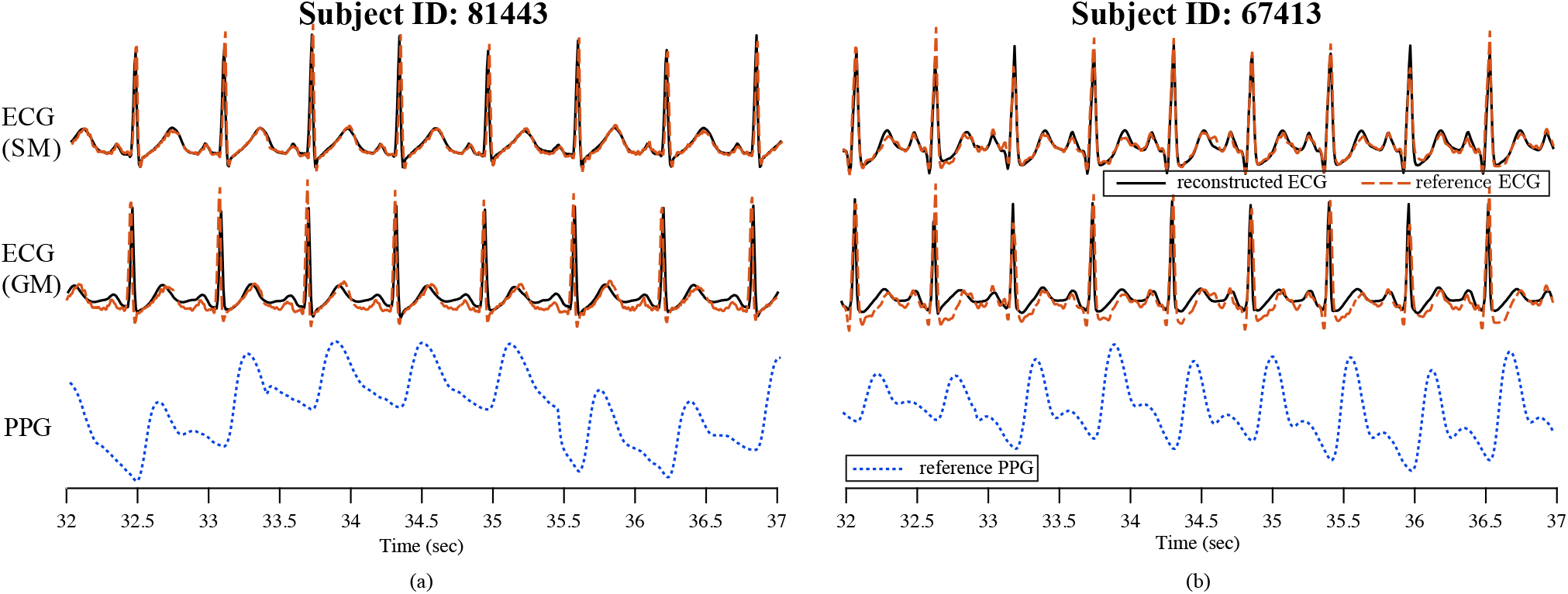
Two qualitative comparisons between the reconstructed ECG signals tested in the subject-specific model (1st row) and group model (2nd row) setup from the MIMIC-III database. (a) The subject is male, 54 years old, and with upper gastrointestinal bleeding. The Pearson’s correlation coefficients are 0.969 in the subject-specific model setup and 0.923 in the group model setup. (b) The subject is male, 52 years old, and with congestive heart failure. The correlation coefficients are 0.959 in the subject-specific model setup and 0.881 in the group model setup.

**TABLE II.**
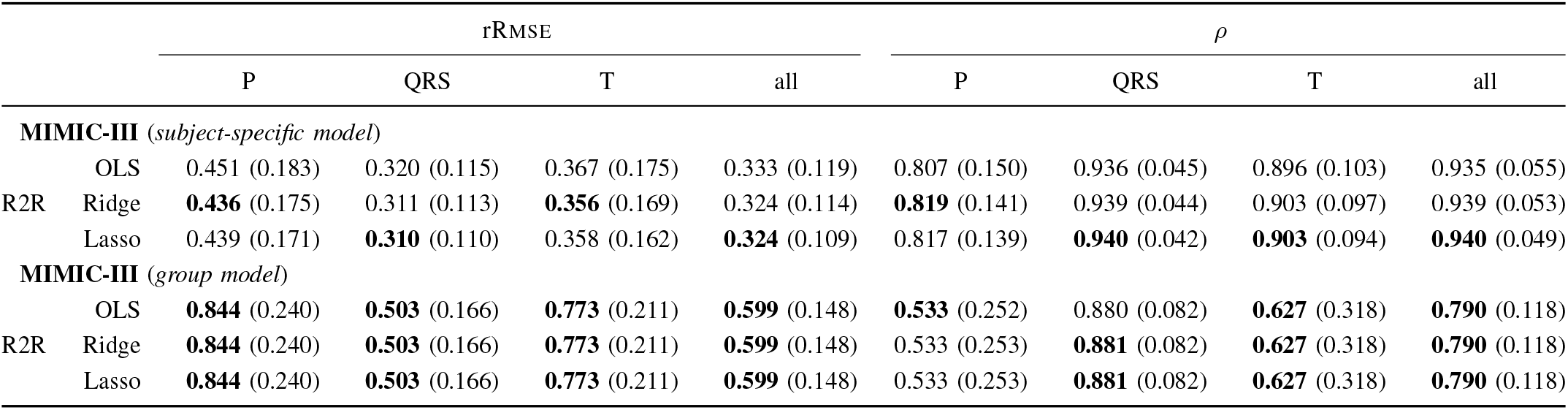
The system performance in test set of the MIMIC-III database in terms of sample mean and standard deviation (in parenthesis) of rRMSE and *ρ*. R2R segmentation using difference combinations of training setup (subject-specific model/group model) and linear regression methods (OLS//LASSO). The best performed entry in each column and training setup is bolded for better visualization.

In Fig. IV-B, we show two five-second segments of the reconstructed ECG waveform in the test set from two subjects using the R2R cycle segmentation scheme with *L_x_* = 18 in the group model setup and *L_x_* = 12 in the subject-specific model setup. The first subject is a 54-year-old male with upper gastrointestinal bleeding, and the second subject is a 52-year-old male with congestive heart failure. We see from the plots that the system retains the major shape of the original ECG waveform except for the P waves of the first subject and s waves of the second subject in the group model setup. The system almost perfectly reconstructs the shape of the ECG waveform in the subject-specific model setup.

In addition to quantitative analysis of the reconstruction performance by Pearson correlation and rRMsE, we also executed a disease classification experiment on the reconstructed ECG signals to show the potential of our proposed method in applications within biomedical health informatics.

First, from the collected MIMIC-III database, we selected 28 patients with five types of cardiac diseases, including congestive heart failure, sT-segment elevated myocardial infarction, non-sT segment elevated myocardial infarction, hypotension, and coronary artery disease. For each patient, we performed the subject-specific model setup ECG reconstruction experiment to obtain the reconstructed ECG signals. The training data is composed of 70 % from the original ECG signals, and the testing data constitutes of the rest 30 % from original ECG signals and all of the reconstructed ECG signals. The detailed distribution of training and testing data concerning disease types is shown in Table III.

**TABLE III.**
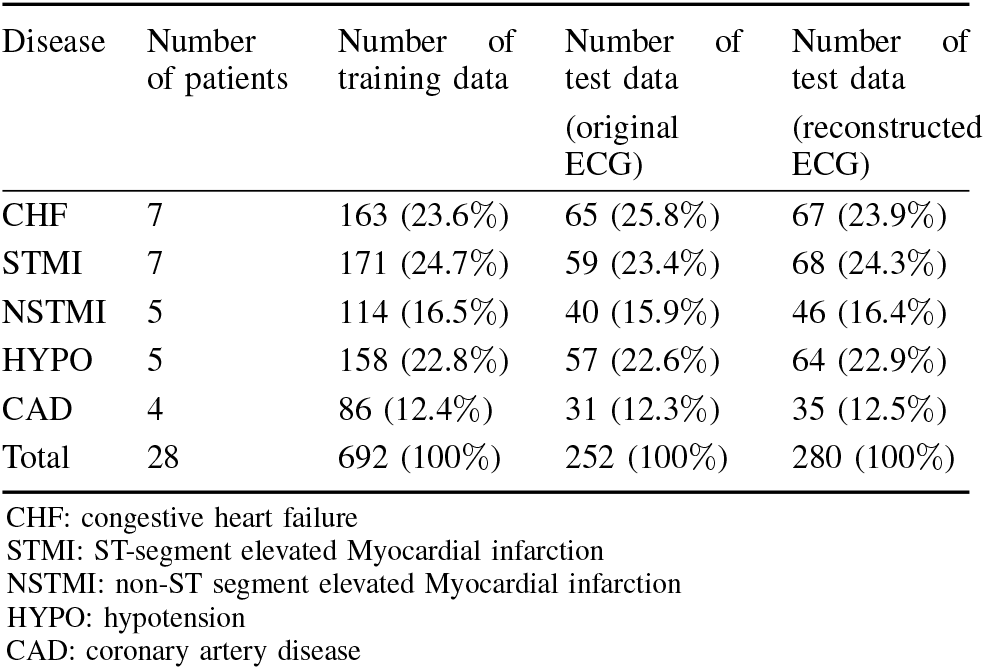
Distribution of training and testing data for disease classification in the MIMIC-III dataset

We applied PCA for dimensionality reduction and sVM classifier with polynomial kernel from sVM library [34]. The confusion matrices for classification are illustrated in Fig. 12 with the reduced dimension as 100. Comparing Figs. 12(a) and 12(b), we conclude that our reconstructed ECG has a comparable classification performance as the original ECG signals. We also include the confusion matrix for the original PPG classification in Fig. 12(c) for reference. The superior performance of classification from the reconstructed ECG signals compared to that of the original PPG signal indicates the fidelity of the reconstructed ECG recordings in the presence of cardiac pathologies.

**Fig. 12.**
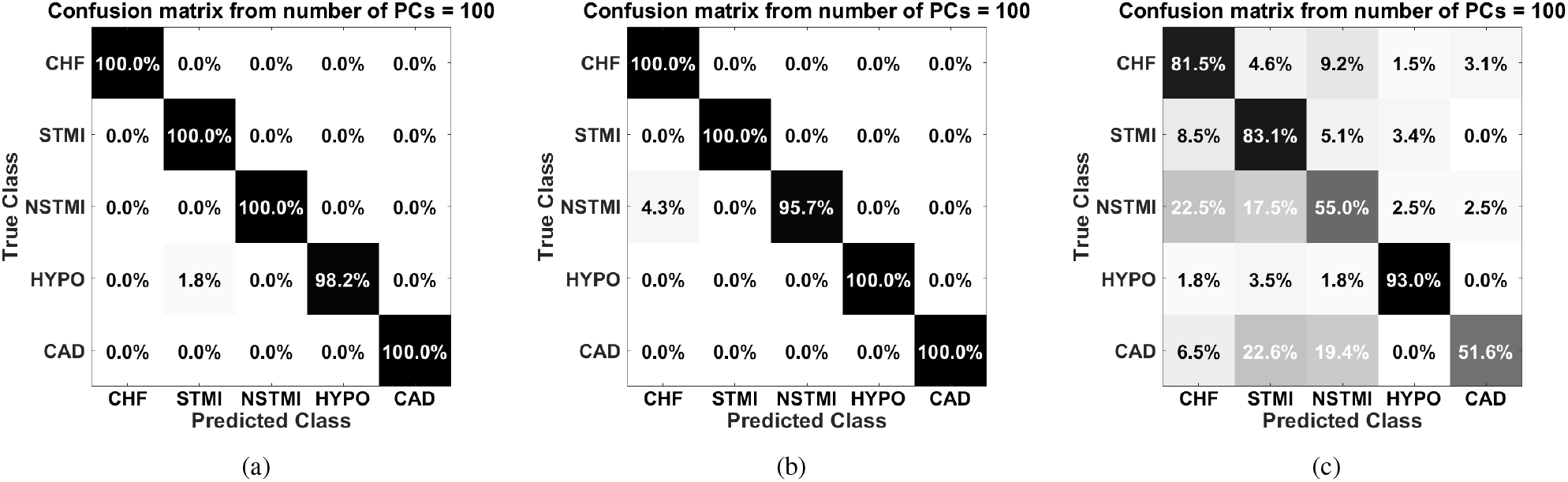
Confusion matrices for classification results using kernel SVM on three types of data: (a) original ECG, (b) inferred ECG, and (c) original PPG.

### C. Experiment 3: Self-collected UMD Dataset

Next, we test the temporal consistency of the proposed system with the self-collected data using consumer-grade sensors. Two subjects participated with informed consent in a physiological data collection under the protocol of #1376735-1 approved by the University of Maryland Institutional Review Board (IRB). Both of them are Asian: a male of 31 years old and a female of 23 years old. According to the best knowledge of the known subjects, none of them have any CVDs or related illnesses of concern. We recorded six 5-min sessions for the first subject and seven sessions for the second subject at different times over two-week. In each session, the subject was asked to wear two devices, namely, EMAY FDA-clear handheld single-lead ECG monitor (Model: EMG-10), and CONTEC pulse oximeter (Model: CMS50E) to record their lead I bipolar ECG signals^4^ and finger-tip PPG signals simultaneously. We asked the subject to wear the PPG sensor on his/her index finger of the right hand and attach the electrodes of the ECG sensor to the palm of the left hand and the back of the right hand. The subjects were asked to sit in front of a table and put their arms on the table as motionless and peacefully as possible to reduce the motion-induced artifacts during the recording time. The sampling rates of the ECG and PPG sensors are 150 and 60 Hz, respectively. We up-sampled both signals to 300 Hz via the bilinear interpolation for consistency consideration and properly aligned the pair of signals.

**TABLE IV.**
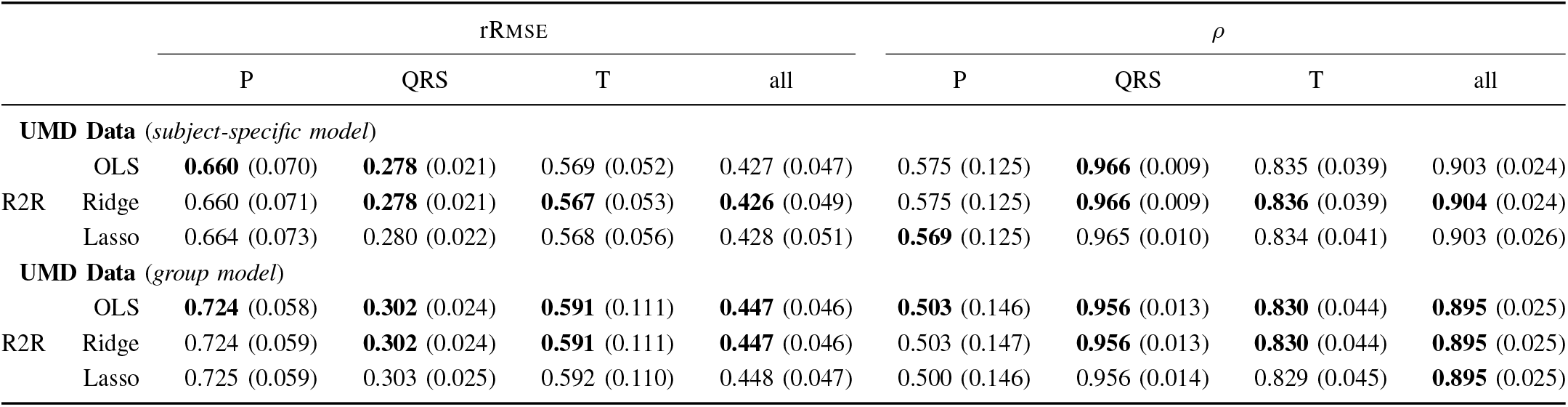
The system performance in test set of the UMD database in terms of sample mean and standard deviation (in parenthesis) of rRMSE and *ρ*. The best performed entry in each column and training setup is bolded for better visualization.

We evaluate the system performance in the subject-specific model and the group model setup. Note that in the subject-specific model setup, the sessions of each subject were first listed chronologically, and **F**\* was trained on the first 80% of the sessions and was tested on the rest of the sessions in order to maximize the temporal difference of the training and test set.

In this experiment, we use the R2R segmentation scheme and set *L_x_* = 12 in subject-specific model setup and *L_x_* = 18 in group model setup. The cycle segmentation process is guided by the peak detection algorithms introduced in [33]. The two algorithms are deployed to detect the R peak of ECG and the onset point of the PPG signal, respectively. Fig. 13 shows one example of the reconstructed waveforms from the 6th session of the first subject. Note that this session is recorded more than one week after the other sessions. From the qualitative result in 2nd and 3rd rows of Fig. 13, we notice that the reconstructed signals match well with the reference ECG in all waves in the condition of long temporal separation from the training set.

**Fig. 13.**
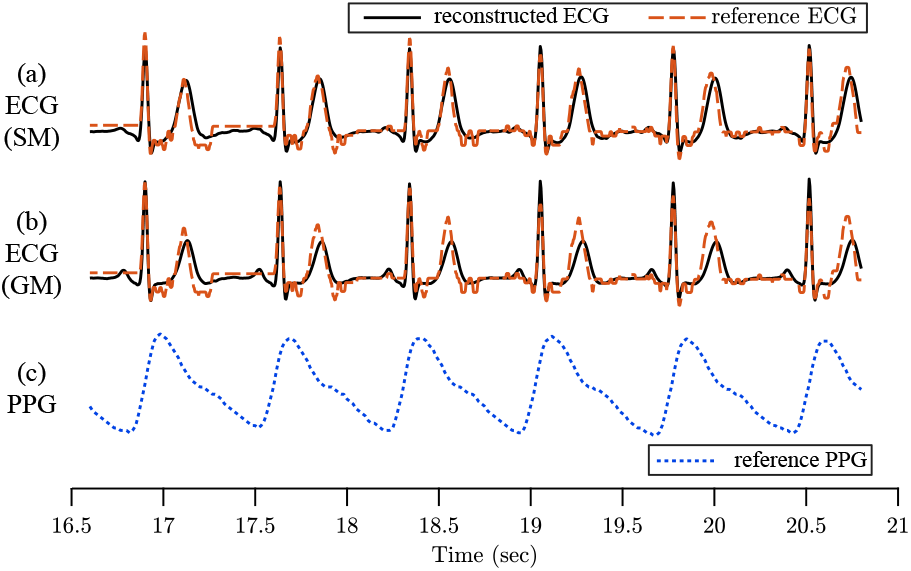
A qualitative comparison among the reconstructed ECG signals tested in (a) subject-specific model and (b) group model setups respectively, from the 6th session of the first subject in the self-collected database. In (a-b), the black line indicates the reconstructed ECG and the orange dashed line refers to the reference ECG. The Pearson’s correlation coefficients for these three cases are 0.917 in the subject-specific model and 0.869 in group model. (c): the corresponding PPG waveform.

Similar to the previous two experiments, we summarized the average performance in different combinations of training setups and regression methods and evaluate each combination in terms of rRMSE and *ρ* in P, QRS, T waves respectively. Notice that in general, the system performs better in subject-specific model compared with the group model. Again, this difference may suggest possible subject-wise difference of the model parameter *b*(*t*), **H**, or α. Consistent observations in this dataset also include better performance in T wave than P wave, and our conjecture remains with the one claimed in Section IV-A.

## V. Discussions

### A. Cycle Segmentation via PPG

We have evaluated the system in Section IV assuming the availability of the ground truth cardiac cycle information obtained from the ECG signal. We now examine a more practical setting when the cycles are estimated solely from the PPG signal, thereby accounting for the real-world constraint that the reference cycle information is unavailable.

The MIMIC-III database introduced in Section IV-B was adopted in this experiment. We segmented the signal according to the onset points of the PPG signal, considering the onset point represents one of the most distinct features within the PPG cycle. We name this segmentation scheme *O2O*.

To single out the contribution to the reconstruction error due to the discrepancy in the waveform shape rather than the misalignment of the ECG peaks, we evaluate O2O after each reconstructed cycle was post-processed to align with the original ECG signal. This was done by shifting each reconstructed ECG cycle in time so that the original and reconstructed ECG signals were matched according to their R peaks. We list the performance metrics in the subject-specific model and group model setups and compare the results with the R2R segmentation in Table V. Note that *ρ* = 0.510 when using O2O segmentation without the peak alignment in the subject-specific model setup, and *ρ* increases to 0.823 once the peak is aligned. The performance statistics reveal that the shape of the waveform is inferred well, and increased error in reconstruction by O2O compared with R2R is mainly due to the misalignment of the signal that has a sample mean and standard deviation of 0.38% and 3.98% in relative cycle length, respectively. This observation is consistent across the group model and the subject-specific model training setups.

**TABLE V.**
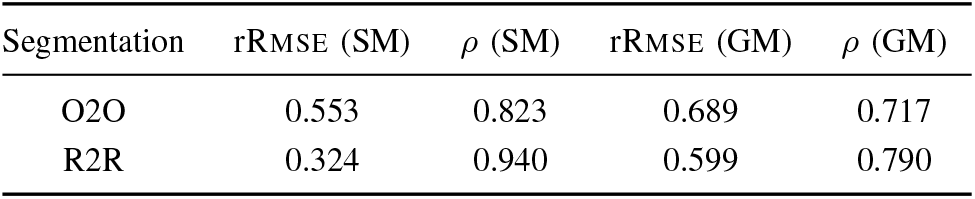
Performance comparison using O2O and R2R cycle segmentation schemes on the MIMIC-III test dataset.

The disease classification experiment was conducted using the O2O segmentation without the peak alignment. We observed a comparable classification accuracy of the reconstructed ECG signal compared with the result when the model was trained with the R2R segmentation. This observation indicates that the ECG reconstruction deviation does not affect the diagnostic power of the reconstructed ECG signal.

### B. Limitations and Extensions of the Proposed Methodology

For some subjects with cardiac complications that influence the morphology of ECG waves, our proposed model in Section II and the corresponding methodology using DCT representations have limitations and may not be able to always faithfully reproduce the ECG signals from PPG, especially when the model is trained in the group model setup. Fig. 14 shows three examples of 5-second long reconstructed ECG signals from the MIMIC-III database using a group model that do not fully capture some detailed characteristics of the original ECG signal. Some other cases that may influence the system performance include motion-induced artifacts and loose contact artifacts in PPG recordings under ambulatory conditions. With a more sophisticated training system and the availability of a larger dataset, we expect such limitations can be addressed.

**Fig. 14.**
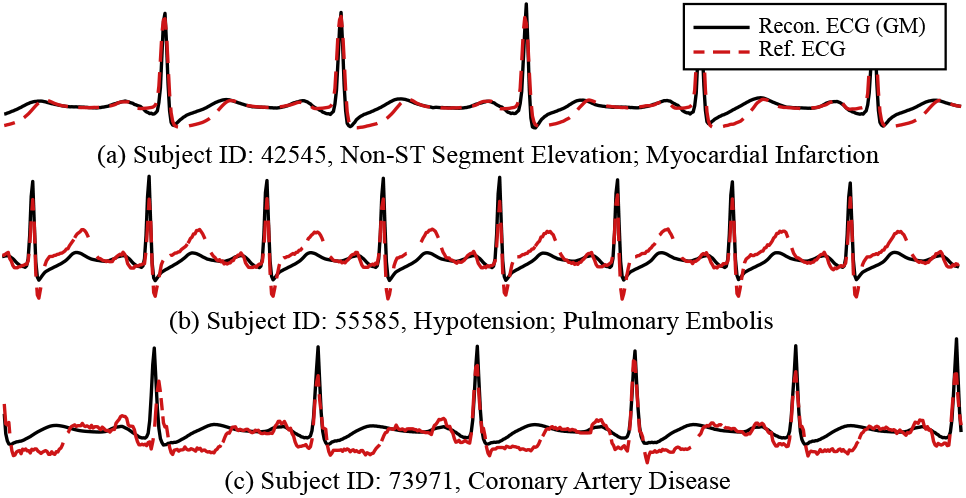
Three examples of the reconstructed ECG signal of low performance in presence of different pathologies of the ECG signal. The reconstructed ECGs fail to capture the waveform during the elevation of the T wave (a, b, c), the T wave (b, c), the P wave (c).

In order to provide more model flexibility in reconstruction, we foresee that the mapping **F** is not limited to a linear transform but can be generalized to nonlinear mappings or transforms (for example, neural networks) and harness more patient data and medical knowledge. Also, the analysis channel of the system is not limited to DCT but can be of other analytical forms, including discrete wavelet transform, discrete Fourier transform, or other parameterized mapping jointly learned with **F** [35]. With further exploration of datasets with detailed profiles of subjects and a larger size of data, a more complex model can be learned based on biomedical, statistical, and physical meanings of the signals to capture the relation of PPG and ECG better. In addition, since ECG is a more adequate and important indicator than PPG for many cardiovascular diseases (CVDs), it has the potential that the developed model, along with the reconstructed ECG, has a significant implication on CVD inference.

While our ongoing follow-up work is to examine these directions beyond the DCT basis to improve the inference accuracy, we recognize that the computational simplicity of linear mappings of DCT coefficients may be suitable for such use cases and health IoT applications that prefer low-cost and low-power system design.

Note that our proposed learning method is not limited to the relation between PPG and single-lead ECG, even though we have focused on the reconstruction of single-lead ECG signals in this paper. With a further effort to collect as rich ECG information as possible, such as 12-lead data, the framework we proposed in this paper can be adapted to gain insights on the relation between PPG and signals of different ECG leads and the feasibility of robust inference for multilead cases. From this angle, our proposed methodology has the potential to offer multi-lead ECG readings from PPG sensors, in addition to the advantage of not requiring active user participation when compared with Apple Watch’s ECG solution.

### C. PPG Denoising via Adaptive Motion Removal

So far, we have discussed ECG reconstruction from clean PPG signals. In practice, PPG signals are prone to motion artifacts. In this section, we show that by leveraging multimodal sensing signals commonly seen in wearable IoT devices and using appropriate adaptive filtering algorithms, we can clean the motion-contaminated PPG signal for robust heart rate (HR) estimation.

As proofs-of-concept, we consider two types of adaptive filters, namely, the normalized least mean squares (NLMS) and the recursive least squares (RLS) adaptive filters [36], [37] for motion removal. We implemented both methods separately and compared the results. Here, we show the result by the RLS method as its overall performance is slightly better than that of the NLMS method. The contaminated PPG is seen as the sum of the underlying cleaned PPG and the noise introduced by motion. Since the three-axis accelerometry signals are correlated with the motion, we use the accelerometry signals to estimate and remove the motion artifacts in PPG. The short-time Fourier-transform is then applied to the clean signals, and the resulting spectrogram is used for HR frequency estimation. We adopt a robust heart rate tracking algorithm [38], which searches for the most probable path by dynamic programming.

The dataset for evaluation is composed of five subjects with 22 sessions of various activities, including running, jumping, rowing, walking, and resting. During the experiment, each subject wears the Empatica E4 watch [39] to collect PPG, accelerometer, and HR data. The subject also wears a Polar H7 chest-strap HR monitor that is considered the gold standard for HR in sports and fitness. We use the data from the built-in PPG sensor and the accelerometers of the E4 watch to track the heart rate. We then compare our approach with the HR recordings directly obtained from the E4 watch using the chest strap as ground truth.

Fig. 15 shows the HR estimation results for a female subject while running. By comparing Fig. 15(d) and Fig. 15(e), we observe that the adaptive filtering has removed most of the motion signals. With our robust trace tracking algorithm, we observe in Fig. 15(g) that it can achieve a more accurate HR tracking result compared to that of the E4 watch under large motion. The average absolute error with respect to the reference HR for our proposed method and the E4 watch are 1.09 bpm and 7.57 bpm, respectively.

**Fig. 15.**
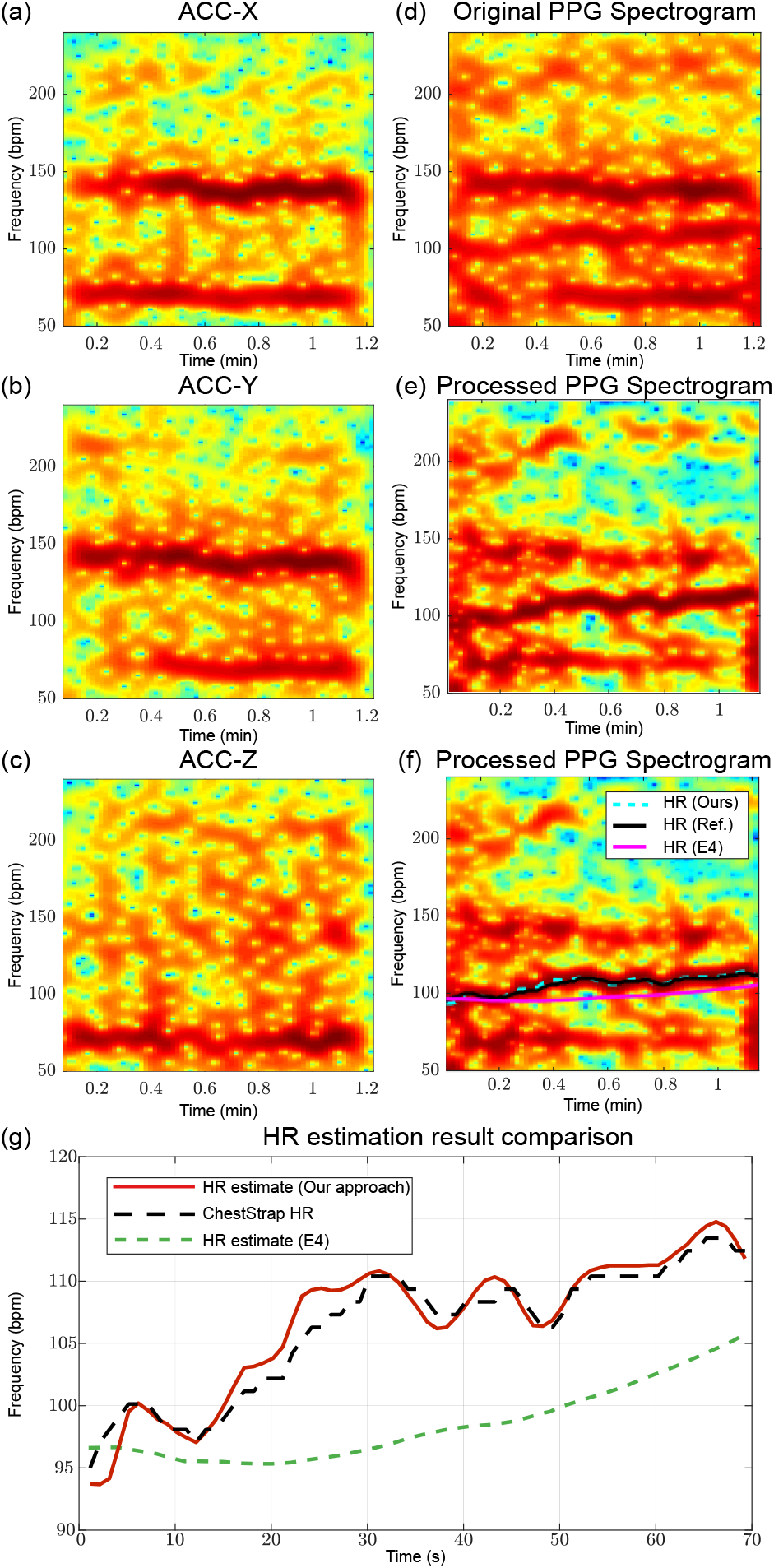
An example from a female subject while running. (a)(b)(c) are the spectrograms for the three-axis accelerometer signals, (d) shows the spectrogram for the original PPG signal, (e) is the spectrogram for the PPG signal after motion artifacts removal, (f) is (e) with HR traces labeled out, and (g) is a zoomed-in version for the HR traces shown in (f).

Fig. 16 shows the overall performance comparison for HR estimation with reference to the chest strap between our proposed smart multimodal filtering method and the HR output by the E4 watch. Fig. 16(a) shows that our approach achieves a high Pearson correlation, 0.954, between HR estimation and the ground truth. In contrast, the heart rate given by E4 is quite off under the strong motion activities with a correlation of 0.657. The Bland-Altman plot in Fig. 16(b) for our approach shows that the bias is just 0.47 bpm whereas that of the E4 HR estimation is 6.2 bpm. The limits of agreement, which is computed as bias ±1.96 times the sample standard deviation of the differences between the estimated HR and the ground truth [40], is [−12.01,12.95] bpm for our method while that of E4 is [−26.66, 39.06] bpm.

**Fig. 16.**
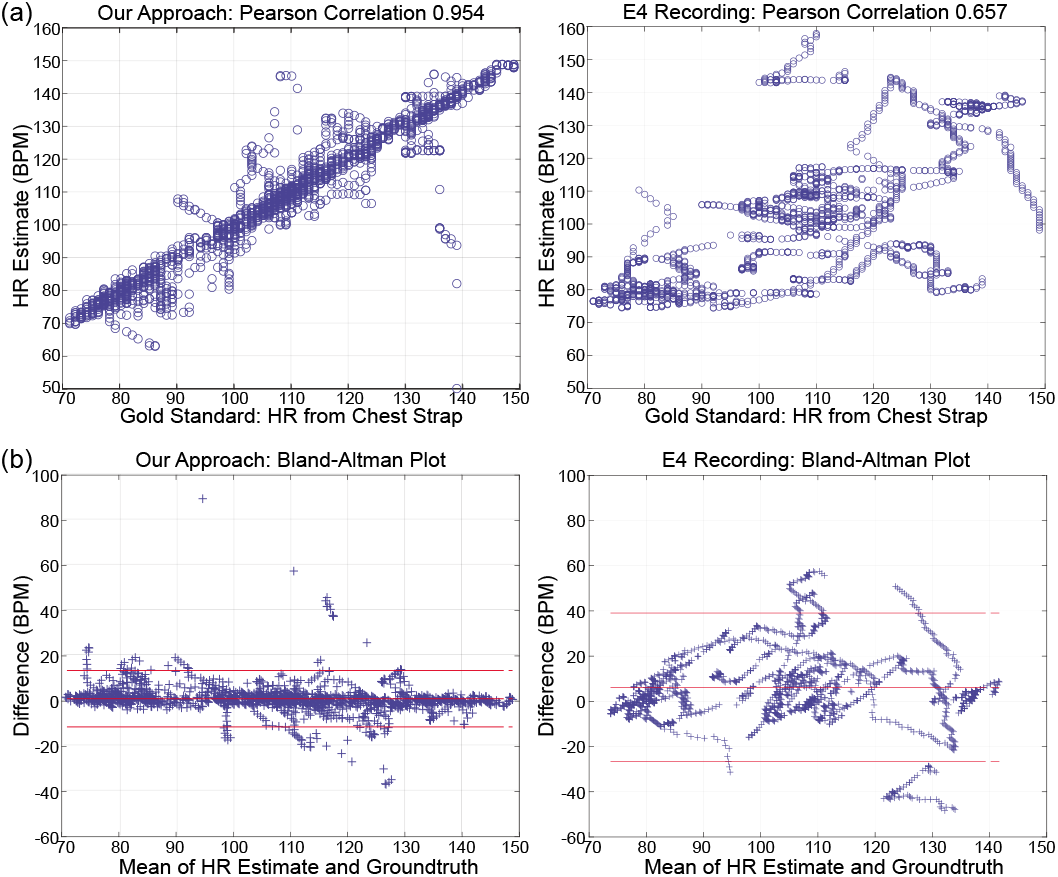
(a) Pearson correlation and (b) Bland-Altman plot w.r.t chest strap reference heart rate for HR estimation results from our approach and E4 watch. The middle, top, and bottom red lines in (b) indicate the averaged difference and the averaged difference ±1.96 times the sample standard deviation of the differences between the estimated HR and the ground truth.

## VI. Conclusion

This paper has presented a learning-based approach to reconstruct ECG signals from PPG signals based on the synergy of a physical model, biomedical knowledge, and data. The algorithm was successfully evaluated with both the subject-specific model and the group model setups on two widely-adopted databases as well as a self-collected database. We have cross-validated the system’s hyperparameters, tested the CVD diagnosis performance using the reconstructed ECG signal and verified the algorithm’s accuracy and consistency at a fine ECG waveform level. As a pilot study, this work demonstrates that with a signal processing and learning system that is designed synergistically, we can reconstruct ECG signals from the more easily obtainable PPG data by exploiting the relation of these two types of cardiovascular related measurements. The DCT domain linear mapping examined in this paper as a proof of concept has potential IoT applications for health and fitness with low-cost, low-power wearable devices, and the proposed framework can be extended to incorporate more sophisticated learning models.

---

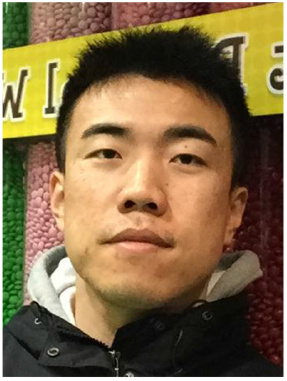

**Qiang Zhu** (S’17–M’20) received his B.E. degree in control science and engineering from Zhejiang University, Hangzhou, China, in 2010, his M.S. degree in control science and engineering from Shanghai Jiao Tong University, Shanghai, China, in 2014, and his Ph.D. degree in electrical engineering from the University of Maryland, College Park, USA, in 2020. He is a research scientist at Facebook since 2020. His research interests are signal processing, machine learning, and information retrieval. He received the Distinguished Teaching Assistance Award in 2016 from the University of Maryland.

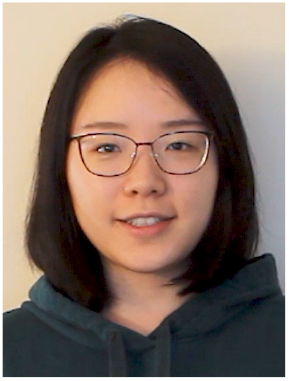

**Xin Tian** (S’18) received the B.E. degree in Optoelectronic Information Science and Engineering from Huazhong University of Science and Technology of China in 2017. She is currently pursuing the Ph.D. degree in electrical and computer engineering at the University of Maryland, College Park. Her current research interests are signal processing, data science, and machine learning in smart health. She was a summer intern at Facebook in 2020. She was selected as a Future Faculty Fellow by the University of Maryland Clark School of Engineering in 2020.

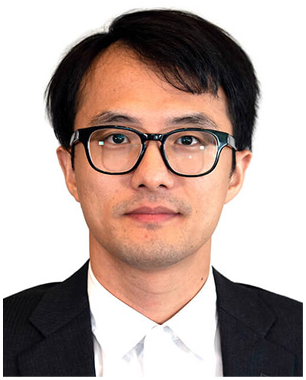

**Chau-Wai Wong** (S’05–M’16) received his B.Eng. and M.Phil. degrees in electronic and information engineering from The Hong Kong Polytechnic University in 2008 and 2010, and the Ph.D. degree in electrical engineering from the University of Maryland, College Park in 2017. He is currently an Assistant Professor at the Department of Electrical and Computer Engineering and the Forensic Sciences Cluster, North Carolina State University. He was a data scientist at Origin Wireless, Inc., Greenbelt, Maryland. His research interests include multimedia forensics, statistical signal processing, machine learning, data analytics, and video coding. Dr. Wong received a Top-Four Student Paper Award, Future Faculty Fellowship, HSBC Scholarship, and Hitachi Scholarship. He was the General Secretary of the IEEE PolyU Student Branch from 2006 to 2007. He was involved in organizing the third edition of the IEEE Signal Processing Cup in 2016 on electric network frequency forensics.

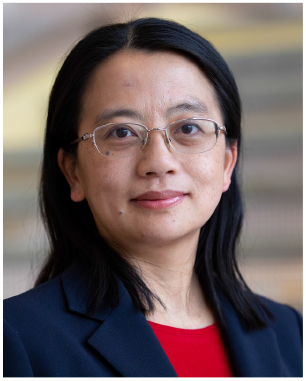

**Min Wu** (S’95–M’01–SM’06–F’11) is a Professor and Associate Dean of Engineering and a Distinguished Scholar-Teacher at the University of Maryland, College Park. She received her undergraduate degrees in engineering and economics with the highest honors from Tsinghua University, Beijing, China, in 1996, and her Ph.D. degree in electrical engineering from Princeton University in 2001. Her research interests include information security and forensics, multimedia signal processing, and applications of data science and machine learning in health and IoT. Her research and education have been recognized by a U.S. NSF CAREER award and an ONR Young Investigator Award, a TR100 Young Innovator Award from the MIT Technology Review, an IEEE Harriett B. Rigas Education Award, and an IEEE SP Society Meritorious Service Award. Dr. Wu chaired the IEEE Technical Committee on Information Forensics and Security, and has served as Vice President - Finance of the IEEE Signal Processing Society and Editor-in-Chief of the IEEE Signal Processing Magazine. She is a Fellow of the IEEE, AAAS, and the U.S. National Academy of Inventor.

## Footnotes

1 Based on the dichromatic model [26], when PPG sensor works in the *reflective mode*, parameters *τ*_0_ and *τ*_1_ in (4) denote the relative reflective strength of the non-pulsatile components and pulsatile components of tissue, respectively.

2 Note that the recording in this database is of high signal quality. In cases when the signal is corrupted by noise or the subject’s motion artifacts, a denoising process is needed to clean the signal before the preprocessing stage.

3 Based on the ICD-9-CM diagnosis codes, we chose those cardiac diseases under the list of “diseases of the circulatory system”, which is corresponding to 390-459 in the ICD-9-CM diagnosis codes. For those non-cardiac diseases, we selected them from other categories, including “injury and poisoning”, “diseases of the respiratory system”, etc.

4 We measure the lead i ECG signal in this experiment considering the easiest accessibility among all leads using the handheld ECG sensor.

## References

[1] Q. Zhu, X. Tian, C.-W. Wong, and M. Wu, “ECG reconstruction via PPG: A pilot study,” in IEEE EMBS International Conference on Biomedical & Health Informatics (BHI), Chicago, IL, May 2019.

[2] “The Global Burden of Disease: 2019 update,” http://ghdx.healthdata.org/gbd-data-tool, accessed: Mar. 2021.

[3] “Sudden death in young people: Heart problems often blamed,” https://www.mayoclinic.org/diseases-conditions/sudden-cardiac-arrest/in-depth/sudden-death/art-20047571, accessed: Mar. 2021.

[4] X. Fan, Q. Yao, Y. Cai, F. Miao, F. Sun, and Y. Li, “Multiscaled fusion of deep convolutional neural networks for screening atrial fibrillation from single lead short ECG recordings,” IEEE Journal of Biomedical and Health Informatics, vol. 22, no. 6, pp. 1744–1753, Nov. 2018.

[5] W. B. Fye, “A history of the origin, evolution, and impact of electrocardiography,” American Journal of Cardiology, vol. 73, no. 13, pp. 937–949, May 1994.

[6] “Maude adverse event report: Conmed corporation invisatrace adult tape wet gel ECG electrodes invisatrace ECG electrodes.” [Online]. Available: https://www.accessdata.fda.gov/scripts/cdrh/cfdocs/cfMAUDE/detail.cfm?mdrfoi_id=2170635

[7] “Zio XT indications, safety, and warnings,” https://www.irhythmtech.com/zio_xt_precautions.

[8] A. Reisner, P. A. Shaltis, D. McCombie, and H. H. Asada, “Utility of the photoplethysmogram in circulatory monitoring,” Anesthesiology: The Journal of the American Society of Anesthesiologists, vol. 108, no. 5, pp. 950–958, May 2008.

[9] W. Karlen, S. Raman, J. M. Ansermino, and G. A. Dumont, “Multiparameter respiratory rate estimation from the photoplethysmogram,” IEEE Trans. on Bio. Eng., vol. 60, no. 7, pp. 1946–1953, Jul. 2013.

[10] T. Aoyagi and K. Miyasaka, “Pulse oximetry: its invention, contribution to medicine, and future tasks.” Anesthesia and Analgesia, vol. 94, no. 1 Suppl, p. S1, Jan. 2002.

[11] R. Payne, C. Symeonides, D. Webb, and S. Maxwell, “Pulse transit time measured from the ECG: an unreliable marker of beat-to-beat blood pressure,” Journal of Applied Physiology, vol. 100, no. 1, pp. 136–141, Jan. 2006.

[12] J. Allen and A. Murray, “Similarity in bilateral photoplethysmographic peripheral pulse wave characteristics at the ears, thumbs and toes,” Physiological Measurement, vol. 21, no. 3, p. 369, Aug. 2000.

[13] R. D. Labati, E. Muñoz, V. Piuri, R. Sassi, and F. Scotti, “Deep-ECG: Convolutional neural networks for ECG biometric recognition,” Pattern Recognition Letters, vol. 126, pp. 78–85, 2019.

[14] G.-M. Lin and H. H.-S. Lu, “A 12-lead ECG-based system with physiological parameters and machine learning to identify right ventricular hypertrophy in young adults,” IEEE Journal of Translational Engineering in Health and Medicine, vol. 8, pp. 1–10, 2020.

[15] G. Zhang, Z. Mei, Y. Zhang, X. Ma, B. Lo, D. Chen, and Y. Zhang, “A noninvasive blood glucose monitoring system based on smartphone ppg signal processing and machine learning,” IEEE Transactions on Industrial Informatics, vol. 16, no. 11, pp. 7209–7218, 2020.

[16] U. R. Acharya, H. Fujita, S. L. Oh, Y. Hagiwara, J. H. Tan, and M. Adam, “Application of deep convolutional neural network for automated detection of myocardial infarction using ecg signals,” Information Sciences, vol. 415, pp. 190–198, 2017.

[17] U. B. Baloglu, M. Talo, O. Yildirim, R. San Tan, and U. R. Acharya, “Classification of myocardial infarction with multi-lead ecg signals and deep cnn,” Pattern Recognition Letters, vol. 122, pp. 23–30, 2019.

[18] A. Y. Hannun, P. Rajpurkar, M. Haghpanahi, G. H. Tison, C. Bourn, M. P. Turakhia, and A. Y. Ng, “Cardiologist-level arrhythmia detection and classification in ambulatory electrocardiograms using a deep neural network,” Nature Medicine, vol. 25, no. 1, pp. 65–69, 2019.

[19] N. Paradkar and S. R. Chowdhury, “Cardiac arrhythmia detection using photoplethysmography,” in 2017 39th Annual International Conference of the IEEE Engineering in Medicine and Biology Society (EMBC). IEEE, 2017, pp. 113–116.

[20] R. Banerjee, A. Sinha, A. D. Choudhury, and A. Visvanathan, “PhotoECG: Photoplethysmography to estimate ECG parameters,” in IEEE International Conf. on Acoustics, Speech and Signal Proc., Florence, Italy, May 2014, pp. 4404–4408.

[21] A. E. Johnson, T. J. Pollard, L. Shen, H. L. Li-wei, M. Feng, M. Ghassemi, B. Moody, P. Szolovits, L. A. Celi, and R. G. Mark, “MIMIC-III, a freely accessible critical care database,” Scientific Data, vol. 3, p. 160035, 2016.

[22] S. A. Martucci, “Symmetric convolution and the discrete sine and cosine transforms,” IEEE Transactions on Signal Processing, vol. 42, no. 5, pp. 1038–1051, May 1994.

[23] H. GholamHosseini, H. Nazeran, and B. Moran, “ECG compression: evaluation of FFT, DCT, and WT performance,” Australas Phys. Eng. Sci. Med., vol. 21, no. 4, pp. 186–192, Dec. 1998.

[24] J. R. Deller, J. H. L. Hansen, and J. G. Proakis, Discrete-Time Processing of Speech Signals. Wiley-IEEE Press, 2000.

[25] R. R. Anderson and J. A. Parrish, “The optics of human skin,” Journal of Investigative Dermatology, vol. 77, no. 1, pp. 13–19, Jul. 1981.

[26] S. A. Shafer, “Using color to separate reflection components,” Color Research & Application, vol. 10, no. 4, pp. 210–218, Dec. 1985.

[27] X. Tao, H. Gao, X. Shen, J. Wang, and J. Jia, “Scale-recurrent network for deep image deblurring,” in Proceedings of the IEEE Conference on Computer Vision and Pattern Recognition, 2018, pp. 8174–8182.

[28] P. Jax and P. Vary, “On artificial bandwidth extension of telephone speech,” Signal Processing, vol. 83, no. 8, pp. 1707–1719, 2003.

[29] J. Allen, “Photoplethysmography and its application in clinical physiological measurement,” Physiological Measurement, vol. 28, no. 3, p. R1, Feb. 2007.

[30] M. P. Tarvainen, P. O. Ranta-Aho, and P. A. Karjalainen, “An advanced detrending method with application to HRV analysis,” IEEE Transactions on Biomedical Engineering, vol. 49, no. 2, pp. 172–175, 2002.

[31] T. Hastie, R. Tibshirani, and J. Friedman, The Elements of Statistical Learning: Data Mining, Inference, and Prediction, 2nd ed. Springer, New York, 2009.

[32] S. Boyd, N. Parikh, E. Chu, B. Peleato, and J. Eckstein, “Distributed optimization and statistical learning via the alternating direction method of multipliers,” Foundations and Trends^®^ in Machine learning, vol. 3, no. 1, pp. 1–122, Jul. 2011.

[33] J. Pan and W. J. Tompkins, “A real-time QRS detection algorithm,” IEEE Transaction on Biomedical Engineering, vol. 32, no. 3, pp. 230–236, Mar. 1985.

[34] C.-C. Chang and C.-J. Lin, “LIBSVM: A library for support vector machines,” ACM Transactions on Intelligent Systems and Technology (TIST), vol. 2, no. 3, p. 27, 2011.

[35] X. Tian, Q. Zhu, Y. Li, and M. Wu, “Cross-domain joint dictionary learning for ecg reconstruction from ppg,” in IEEE International Conf. on Acoustics, Speech and Signal Proc. Barcelona, Spain: IEEE, 2020, pp. 936–940.

[36] T. Schäck, C. Sledz, M. Muma, and A. M. Zoubir, “A new method for heart rate monitoring during physical exercise using photoplethysmographic signals,” in Signal Processing Conference (EUSIPCO), 2015 23rd European. IEEE, 2015, pp. 2666–2670.

[37] E. Khan, F. Al Hossain, S. Z. Uddin, S. K. Alam, and M. K. Hasan, “A robust heart rate monitoring scheme using photoplethysmographic signals corrupted by intense motion artifacts,” IEEE Transactions on Biomedical Engineering, vol. 63, no. 3, pp. 550–562, 2016.

[38] Q. Zhu, M. Chen, C.-W. Wong, and M. Wu, “Adaptive multi-trace carving for robust frequency tracking in forensic applications,” IEEE Transactions on Information Forensics and Securisty, vol. 16, pp. 1174–1189, Oct. 2020.

[39] M. Garbarino, M. Lai, D. Bender, R. W. Picard, and S. Tognetti, “Empatica E3—A wearable wireless multi-sensor device for real-time computerized biofeedback and data acquisition,” in 2014 4th International Conf. on Wireless Mobi. Comm. and Healthcare-Transforming Healthcare Through Innovations in Mobi. and Wireless Tech. IEEE, 2014, pp. 39–42.

[40] D. Giavarina, “Understanding Bland Altman analysis,” Biochemia medica, vol. 25, no. 2, pp. 141–151, 2015.

